# RORc expressing immune cells negatively regulate tertiary lymphoid structure formation and support their pro-tumorigenic functions

**DOI:** 10.1101/2022.07.02.498540

**Authors:** Einat Cinnamon, Ilan Stein, Elvira Zino, Stav Rabinovich, Yehuda Shovman, Yehuda Schlesinger, Tomer-Meir Salame, Shlomit Reich-Zeliger, Michal Lotem, Yinon Ben-Neriah, Oren Parnas, Eli Pikarsky

**Affiliations:** The Concern Foundation Laboratories at The Lautenberg Center for Immunology and Cancer Research, Israel-Canada Medical Research Institute, Faculty of Medicine, The Hebrew University, Jerusalem, Israel; Department of Pathology, Hadassah-Hebrew University Medical Center, Jerusalem, Israel; Flow Cytometry Unit, Life Sciences Core Facilities, Weizmann Institute of Science, Rehovot, Israel; Department of System Immunology, Weizmann Institute of Science, Rehovot, Israel; Sharett Institute of Oncology, Hadassah-Hebrew University Medical Center, Jerusalem, Israel

**Keywords:** Tertiary lymphoid structure, liver cancer, cholangiocarcinoma, RORc, B cells, CD4 cells, CD8 cells, cell depletion, single cell RNA sequencing, CyTOF

## Abstract

Tertiary lymphoid structures (TLSs) are formed in many cancer types and have been correlated with better prognosis and response to immunotherapy. In liver cancer, TLSs have been reported to be pro-tumorigenic as they harbor tumor progenitor cells and nurture their growth. The processes involved in TLS development and the acquisition of a pro- or anti-tumorigenic phenotype in cancer are largely unknown. RORc expressing immune cells have been previously implicated in TLS formation, however we find that they are not necessary for TLS neogenesis in the context of inflammation-associated liver cancer. On the contrary, RORc expressing cells negatively regulate TLS formation, since in their absence TLSs form in excess. CD4 cells are essential for liver TLS formation whereas B cells are required for TLS formation specifically in the absence of RORc expressing cells. Importantly, in chronically inflamed livers lacking RORc expressing cells, TLSs become anti-tumorigenic, resulting in reduced tumor load. Comparing liver pro- and anti-tumorigenic TLSs by transcriptional, proteomic and immunohistochemical analyses, revealed enrichment of exhausted CD8 cells that retained effector functions as well as germinal center B cells and plasma cells in anti-tumorigenic TLSs. Cell depletion experiments revealed a role mainly for B cells in limiting tumor development, possibly via tumor directed antibodies. Thus, RORc expressing cells negatively regulate B cell responses, and facilitate the pro-tumorigenic functions of hepatic TLSs.

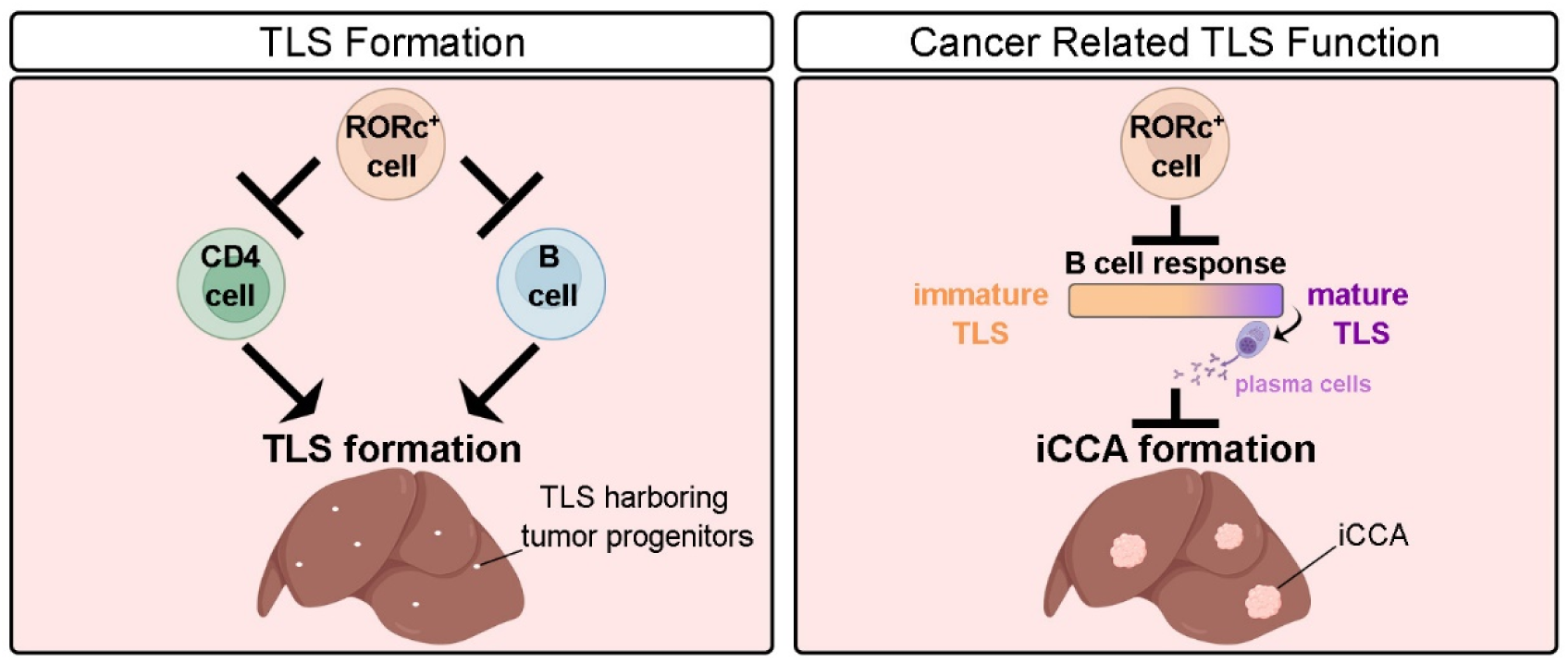

## Introduction

Tertiary lymphoid structures (TLSs) are organized aggregates of immune cells that form in nonlymphoid tissues due to chronic inflammation, autoimmunity and cancer. TLSs have been observed in various cancer types, such as non–small cell lung cancer, colorectal cancer, ovarian cancer and melanoma, and are associated with better patient survival (Sautès-Fridman et al., 2019; Schumacher & Thommen, 2022). However, previous studies have described a pro-tumorigenic role for TLSs in breast, lung and liver cancer (Figenschau et al., 2015; Finkin et al., 2015; Joshi et al., 2015). In a mouse model of liver cancer developing in the background of chronic inflammation, TLSs harboring cancer progenitor cells were found to promote tumor formation. The development of TLSs and tumors depended on an intact adaptive immune system, and TLS-derived Lymphotoxin β was shown to promote cancer progenitor growth (Finkin et al., 2015).

While TLSs in the inflamed liver parenchyma are associated with poor prognosis in human hepatocellular carcinoma (HCC) (Meylan et al., 2020), intratumor TLSs in the liver are associated with a favorable prognosis (Calderaro et al., 2019). Intrahepatic cholangiocarcinoma (iCCA), the second-most common primary liver malignancy after HCC (Bertuccio et al., 2019), showed a similar pattern (Ding et al., 2022). Understanding the neogenesis and phenotypes of TLSs could have important implications for immunological treatments in cancer in general and in liver cancer specifically.

TLS formation was compared to the formation of secondary lymphoid organs (SLO). SLO formation is initiated by the colonization of lymph node anlagen by hematopoietic lymphoid tissue inducer (LTi) cells which are CD4^+^ CD3^−^ CD45^+^ innate lymphoid cells, characterized by the expression of the RORgt (the shorter isoform of RORg, encoded by the RORc locus) and Id2 transcription factors. The interaction between LTi and lymphoid tissue organizer cells leads to recruitment of immune cells to the lymphoid niche (Schumacher & Thommen, 2022). While LTi cells are essential for SLO formation, their relevance for TLS neogenesis has been studied mainly in the context of autoimmunity and chronic infection, and their role in the context of cancer remains to be determined.

RORc, a nuclear hormone receptor, plays a crucial role in the differentiation and development of T helper 17 (Th17) cells, innate lymphoid cells (LTi and ILC3), and γδT cells. Mice lacking RORc fail to develop these cell types and do not form lymph nodes and Peyer’s patches. We asked whether RORc expressing cells (RORcECs) are required for hepatic TLS neogenesis under chronic inflammation leading to liver cancer. Surprisingly, we show that RORcECs have a dual role in TLS-related liver cancer, first by paradoxically suppressing TLS formation and second by contributing to promotion of TLS-dependent liver cancer formation.

## Results

### RORc expressing cells are not necessary for TLS neogenesis but rather negatively regulate their formation

IKKβ(EE)^Hep^ mice express an active form of IKKβ in hepatocytes, mimicking chronic inflammation (Finkin et al., 2015). Few RORc positive cells were observed within liver TLSs in these mice (Fig. 1A, arrowhead, inset). To investigate the role of RORcECs in hepatic TLS formation, IKKβ(EE)^Hep^ mice (denoted IKK) were crossed with whole body RORc knockout mice (RORc^tm2Litt^ KO mice, denoted RORc KO) (Eberl et al., 2004). The resulting IKK RORc KO mice were analyzed alongside mice heterozygous (RORc Het) or homozygous (RORc KO) for the RORc mutant allele, as well as with IKK RORc Het mice (which are phenotypically similar to IKK mice). Since around 50% of RORc KO mice develop T cell lymphomas (Ueda et al., 2002), mice with thymus/spleen enlargement or hepatic lymphoma were excluded from the analysis. Mice were examined under different conditions: A) 6 and 12 months without carcinogen treatment (spontaneous), where the number and size of TLSs containing tumor progenitors in IKK mice increased over time, and tumors appear after 12 months; B) 6 months after a single injection of the carcinogen diethylnitrosamine (DEN), which accelerated TLS and tumor formation. Therefore, 6 months old DEN-injected IKK mice have developed both TLSs harboring tumor progenitors and tumors (Finkin et al., 2015; Tolba et al., 2015).

**Figure 1:**
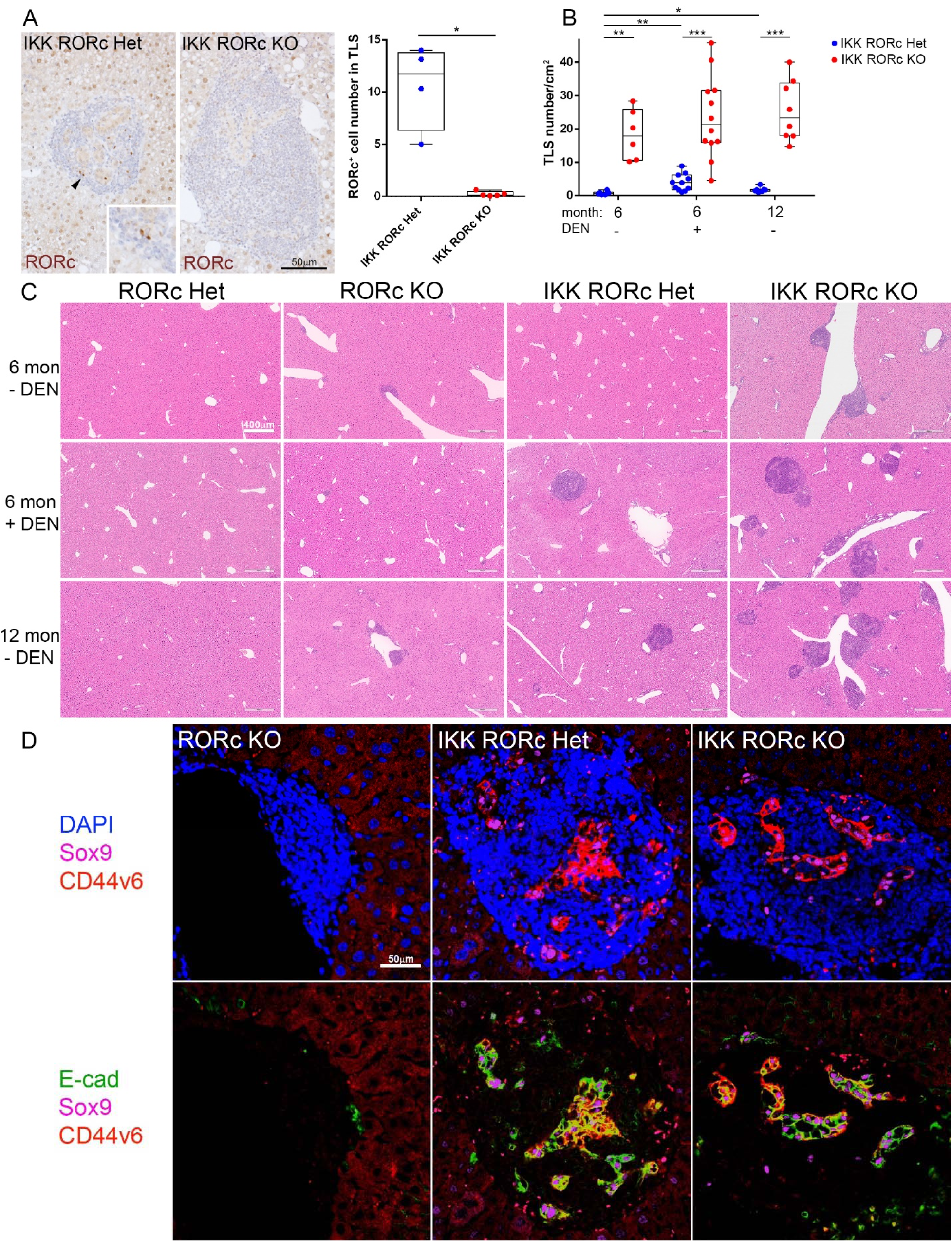
RORc expressing cells are not necessary for liver TLS neogenesis, but rather negatively regulate TLS formation. **(A)** TLSs of DEN-injected 6 months old IKK RORc Het and IKK RORc KO mice immunostained for RORc and the corresponding quantification (each data point is an average of several TLSs. For all figures, each data point represents a single mouse). Arrowhead: RORc expressing cells (also shown in the inset). **(B)** Numbers of TLSs harboring tumor progenitors. **(C)** H&E stained images of liver sections in spontaneous conditions at 6 months (6 mon - DEN), 6 months following DEN injection (6 mon + DEN) and spontaneous conditions at 12 months of age (12 mon - DEN). **(D)** Triple immunofluorescent staining for Sox9, CD44v6 and E-cadherin. DAPI (blue) stains nuclei and delineates the TLSs. Sox9 (pink, nuclear) and CD44v6 (red, membranous) are tumor progenitor markers. E-cadherin (green) marks epithelial cells. Note that TLSs in RORc KO mice do not harbor tumor progenitor cells. Statistical tests applied: Mann-Whitney (A); Kruskal-Wallis with Bonferroni correction (B).

Surprisingly, spontaneous and DEN-injected IKK RORc KO mice retained the ability to develop TLSs harboring tumor progenitors, similar to IKK RORc Het mice, suggesting that RORcECs are not essential for TLS development. Instead, the abundance of TLSs with tumor progenitors was 5- to 27-folds higher in IKK RORc KO mice compared to IKK RORc Het mice in DEN-injected mice as well as spontaneous conditions (Fig. 1B,C, Fig. S1A and Fig. S2). In all conditions, IKK RORc KO mice developed excess TLSs that were smaller than 400μm in diameter (Fig. S1C). Tumor progenitors within TLSs were identifiable by the expression of Sox9 and CD44v6 and through H&E staining (Fig. 1D and Fig. S1A). As expected, RORcECs were absent in TLSs from IKK RORc KO mice (Fig. 1A). In both IKK RORc Het and IKK RORc KO mice, TLSs increased in size over time, especially with DEN treatment or aging (12 months). However, spontaneous IKK RORc KO mice at 6 months only developed small-sized TLSs (below 200μm) (Fig. S1C), suggesting that RORcECs primarily inhibit TLS neogenesis rather than increase in size. These results suggest that RORcECs are not necessary for TLS neogenesis, but rather negatively regulate TLS formation.

Interestingly, in IKK RORc Het mice, DEN administration or aging led to increased TLSs with tumor progenitors. In contrast, neither DEN nor aging contributed to further TLS formation in IKK RORc KO mice (Fig. 1B). The inability of increased antigen load to further increase TLS numbers in the absence of RORcECs suggests that the quantity of TLSs that can form in the liver is restrained.

Spontaneous and DEN-injected RORc KO mice and IKK RORc KO mice also developed small TLSs (less than 150μm in diameter) lacking tumor progenitor cells (Wichner et al., 2013; see ‘image analysis’ in methods) which increased with time or DEN treatment (Fig. S1B). These were 2-4 times less abundant than TLSs harboring tumor progenitors and were not apparent in IKK RORc Het mice (Fig. 1C,D and Fig. S1B).

In conclusion, RORcECs are not necessary for TLS neogenesis but instead negatively regulate TLS formation in the liver.

### CD4 cells are essential for liver TLS formation whereas B cells are required for TLS formation specifically in the absence of RORc

Next, we investigated the role of adaptive immune cells in TLS formation by depleting CD4 (αCD4), CD8 (αCD8), or B cells (αCD20). Initially, we assessed the requirement of these cell types for TLS maintenance in DEN-injected IKK RORc Het mice, where TLSs start to form at around 3 months (Finkin et al., 2015). αCD8 and αCD20 depleting antibodies (from mouse host) were given after TLS started to form (4.5 to 6.5 months), while αCD4 antibodies (from rat host) were given for a shorter period (5.5 to 6.5 months) to avoid a humoral response against the depleting antibodies (Fig. 2A). CD8, CD4 and B cells occupy an average of 5.4%, 7.5% and 24.5% of the TLS area, respectively. Efficient depletion was observed in the spleen and liver (Fig. S3A-D, and Fig. S4A-D). CD4 depletion, but not CD8 or CD20 depletion, resulted in decreased TLS numbers (both below and above 400μm in diameter) (Fig. 2B-D and Fig. S5), suggesting that CD4 cells are necessary for TLS maintenance.

**Figure 2:**
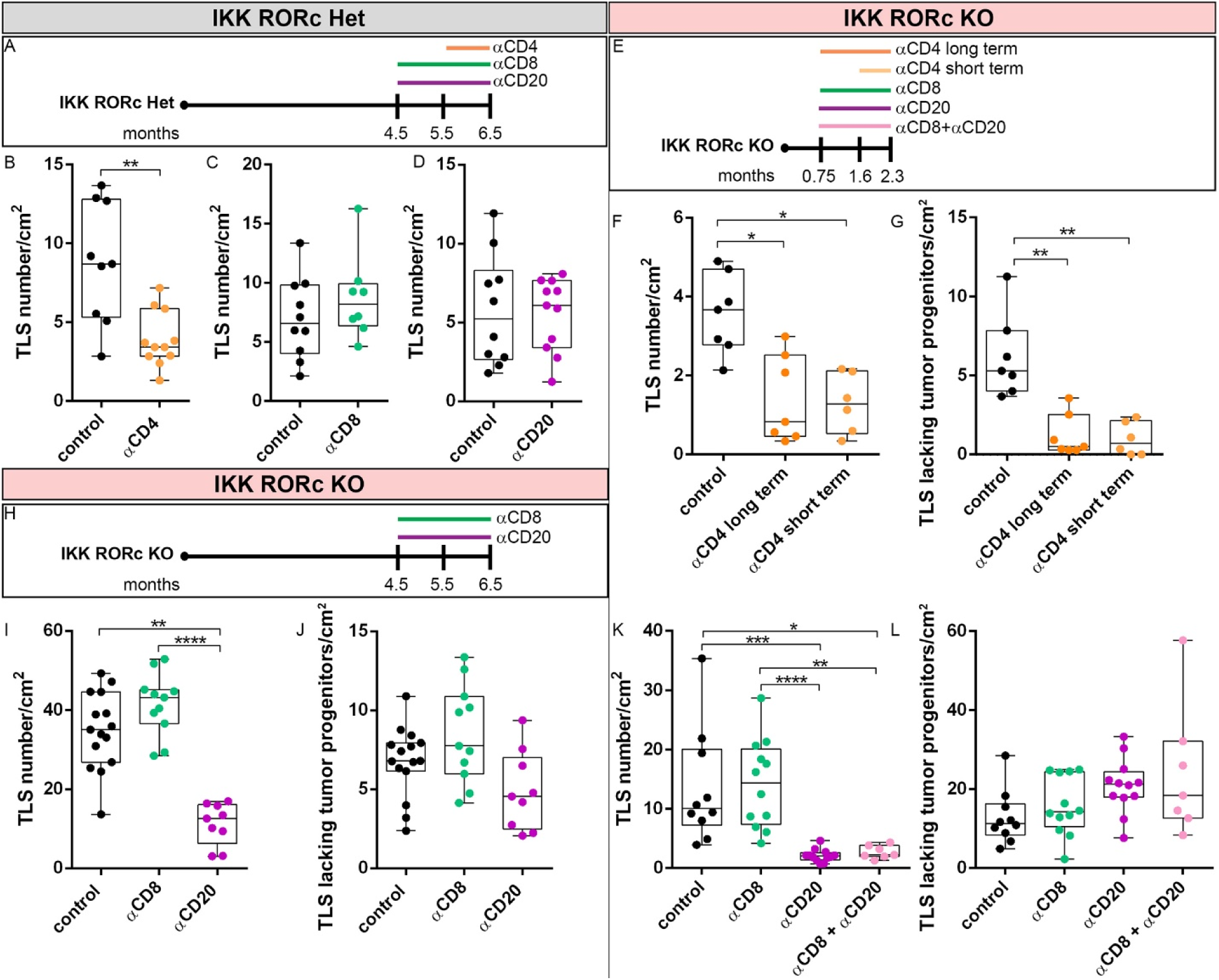
CD4 and B cells are required for TLS formation. **(A)** Antibody depletion regime in DEN-injected IKK RORc Het mice. Cell depletion treatments began after TLSs started to form. αCD4 depleting antibodies were given between 5.5 to 6.5 months, while αCD8 and αCD20 (B cells) depleting antibodies were given between 4.5 to 6.5 months. Mice were sacrificed at 6.5 months. Numbers of TLSs harboring tumor progenitors following CD4 depletion **(B)**, CD8 depletion **(C)** and CD20 depletion **(D)** in IKK RORc Het mice, as depicted in 2A. **(E)** Antibody depletion regime in DEN-injected IKK RORc KO mice. Cell depletion treatments began at 3 weeks, before TLSs started to form [αCD4 long term, αCD8, αCD20 or their combination (αCD8+αCD20)]. αCD4 treatment (short term) began at 1.6 months, after TLSs started to form. Mice were sacrificed at 2.3 months. Numbers of TLS harboring **(F)** and lacking **(G)** tumor progenitors following CD4 depletion as depicted in 2E. **(H)** Antibody depletion regime in DEN-injected IKK RORc KO mice. Cell depletion treatments began after TLSs started to form. αCD8 and αCD20 depleting antibodies were given between 4.5 to 6.5 months. Number of TLSs harboring **(I)** and lacking **(J)** tumor progenitors following CD8 and CD20 depletion as depicted in 2H. Numbers of TLSs harboring **(K)** and lacking **(L)** tumor progenitors following CD8, CD20 or combined CD8+CD20 depletion in IKK RORc KO mice as depicted in 2E. Statistical tests applied: Mann-Whitney (B-D), Kruskal-Wallis with Bonferroni correction (F,G, I-L).

In DEN-injected IKK RORc KO mice, TLSs are formed excessively and at an accelerated rate [at 3 weeks TLSs have not formed yet but by the age of 1.6 months TLSs are already seen (data not shown)], and at 2.3 months they reach 20-50% of their amount at 6 months (compare control in Fig. 2F,K to 6 months +DEN in Fig. 1B). These rapid dynamics allowed us to test whether CD4 cells are required for TLS neogenesis or maintenance by comparing mice that received αCD4 treatment before or after TLSs have formed (αCD4 long-term, starting at 3 weeks, compared to αCD4 short-term, starting at 1.6 months) (Fig. 2E; for depletion efficiency see Fig. S3E,F and Fig. S6A). Both long- and short-term αCD4 depletion treatments resulted in a similar decrease in the numbers of TLS harboring or lacking tumor progenitors without an additive effect of the long-term treatment (Fig. 2F,G and Fig. S7). These results suggest that CD4 cells are necessary for TLS maintenance, however, a possible role for CD4 cells also in neogenesis cannot be ruled out.

Depletion of CD8 cells in DEN-injected IKK RORC KO mice, after TLSs have already been established (between 4.5 and 6.5 months, Fig. 2H) or before TLS start to form (3 weeks to 2.3 months, Fig. 2E), did not affect the numbers of TLSs harboring tumor progenitors (Fig. 2I,K, Fig. S8A-C, Fig. S9 and Fig. S10; for depletion efficiency see Fig. S3E,G,H,I, Fig. S4E and Fig. S6B) or TLSs lacking tumor progenitors (Fig. 2J,L), as seen in IKK RORc Het mice (Fig. 2C). However, in contrast to IKK RORc Het mice (Fig. 2D), B cell depletion in IKK RORc KO mice, both after TLSs have already been established and before TLS start to form, led to a significant decrease in numbers of TLSs harboring tumor progenitors (Fig. 2I,K, Fig. S8D-F Fig. S9 and Fig. S10; for depletion efficiency see Fig. S3E,G,H,J, Fig. S4F,G and Fig. S6C), indicating the requirement of B cells for TLS formation only in the absence of RORc. TLS numbers were more strongly reduced when B cells were depleted prior to TLS formation compared to after their formation [4.9 fold reduction in 2.3 months old IKK RORc KO mice (Fig. 2K) compared to 2.7 fold reduction in 6.5 months old IKK RORc KO mice, (Fig. 2I)], suggesting that B cells contribute to neogenesis of TLSs rather than their maintenance. However, a possible role for B cell also in TLS maintenance cannot be ruled out. Intriguingly, B cell depletion in IKK RORc KO did not reduce the number of TLSs lacking tumor progenitors, as did CD4 depletion, suggesting that B cells support tumor progenitor development within TLSs, but do not affect immune cell aggregation (Fig. 2J,L and Fig. S10).

Interestingly, B cell depletion prior to TLS formation in IKK RORC KO mice caused a significant three-fold increase in the CD8 population in the spleen and liver (Fig. S3G and data not shown). This expanded CD8 population could potentially target tumor progenitor cells, leading to their reduced survival. To test if the B cell depletion effect is mediated by CD8 cells we depleted both CD8 and B cells (Fig. 2E; see depletion efficiency in Fig. S6B,C). This resulted in reduced formation of TLSs harboring tumor progenitors (Fig. 2K, Fig. S9 and Fig. S10) and had no impact on TLSs lacking tumor progenitors (Fig. 2L and Fig. S10), mirroring the effects of B cell depletion alone. These results indicate that B cells promote the development of tumor progenitors in TLSs independently of CD8 cells.

Overall, our findings indicate that CD4 cells are required for the maintenance of TLSs, while B cells are necessary for TLS neogenesis only in the absence of RORcECs.

### iCCA load is decreased in IKK mice lacking RORc expressing cells

Previously (Finkin et al., 2015) we demonstrated that constitutively activating IKK in hepatocytes induces formation of TLSs which then promote the growth of aggressive iCCAs. Thus, we expected that the increased numbers of TLSs in IKK RORc KO mice will further increase iCCA load. Surprisingly however, DEN-injected IKK RORc KO mice exhibited a 4.7 fold decrease in iCCA formation despite a significant 5.5 fold increase in TLS numbers, compared to IKK RORc Het mice (Fig. 3A,A’,B,D,D’,E,H). The ratio of iCCA number to TLS number, reflecting the ability of TLSs to promote iCCAs, was also significantly reduced by 23.5 fold in IKK RORc KO mice (Fig. 3I). HCC formation remained unaffected (Fig. 3A,A’,C,D,D’,F,G). This suggests a shift in the phenotype of TLSs from pro-to anti-tumorigenic upon deletion of RORc. Thus, RORcECs may control the phenotype of hepatic TLSs. Both RORγ and its shorter isoform, RORγt, are deleted in RORc KO mice. While RORγt is expressed specifically in the immune compartment, RORγ is expressed in many cell types, including hepatocytes (Eberl, 2017). However, deleting RORc specifically in hepatocytes of DEN-injected IKK mice (using a floxed RORc mutant driven by Albumin-Cre, termed IKK RORc^HepKO/HepKO^ mice) did not alter TLS, HCC or iCCA abundance (Fig. 3J). This suggests that hepatic RORc expression does not regulate TLS or liver cancer development. Instead, the elimination of RORc in the immune compartment likely contributes to the increased TLS neogenesis and decreased iCCA formation observed in IKK RORc KO mice. These findings indicate that immune RORcECs play a role in facilitating the pro-tumorigenic functions of hepatic TLSs.

**Figure 3:**
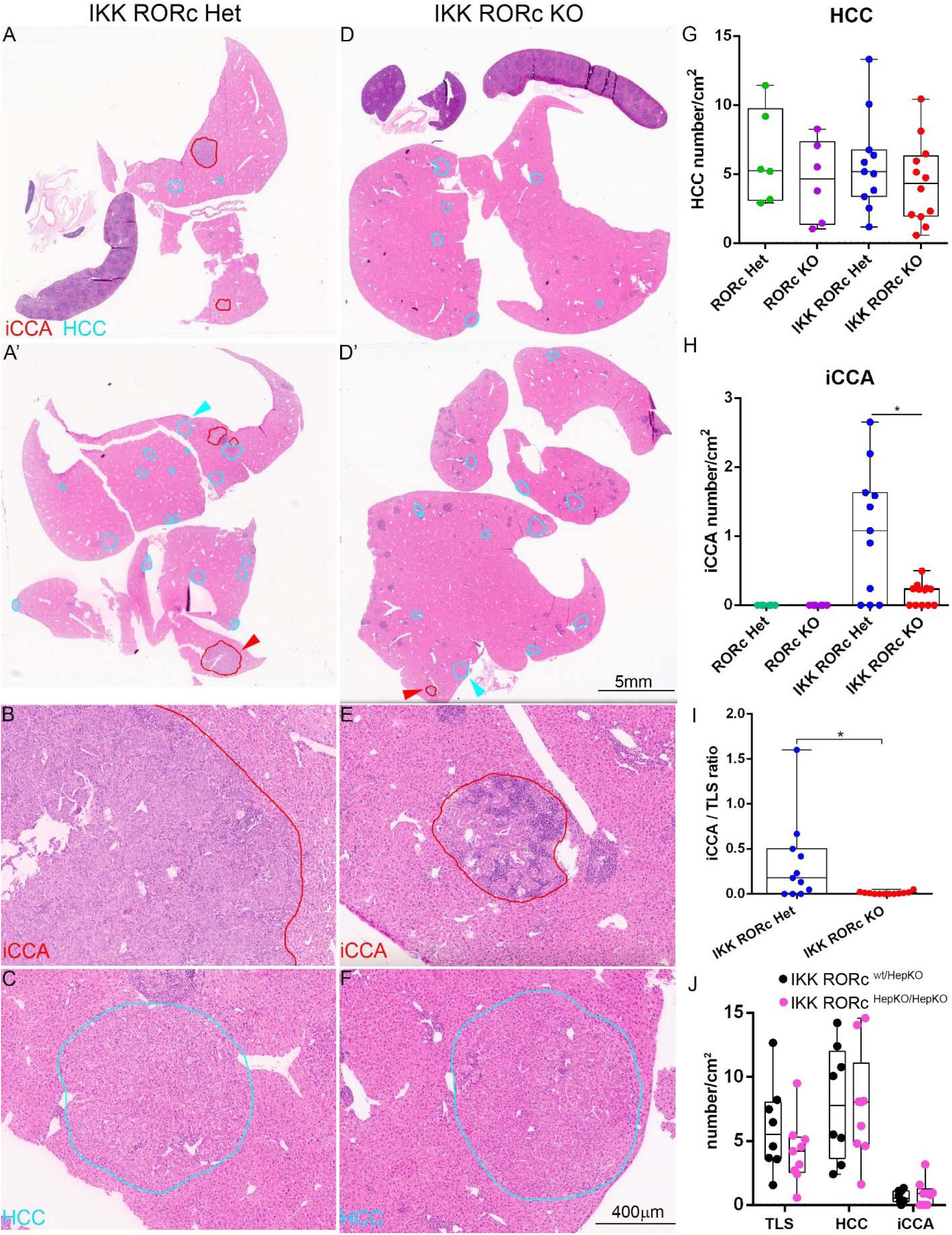
iCCA but not HCC formation is supported by RORc expressing cells. **(A-F)** Low and high magnification images of H&E stained liver sections in DEN-injected 6 months old IKK RORc Het and IKK RORc KO mice. Two sections per mouse, representing the entire liver, are shown **(A, A’** and **D, D’)**. HCCs are circled in light blue and iCCAs in red. Tumors indicated by arrowheads in A’ and D’ are shown in higher magnification in B,C and E,F, respectively. HCC **(G)** and iCCA **(H)** numbers. **(I)** Ratio of iCCA/TLS number. **(J)** TLSs harboring tumor progenitors, HCC and iCCA numbers in DEN-injected 6 months old IKK RORc^wt/HepKO^ and IKK RORc^HepKO/HepKO^ mice. Statistical tests applied: (G) Kruskal-Wallis with Bonferroni correction; (H) Kruskal-Wallis; (I,J) Mann-Whitney.

### CD8 cells with an exhausted/effector phenotype and class-switched B cells are enriched in livers of IKK RORc KO mice

Our findings provide an opportunity for comparing pro- and anti-tumorigenic hepatic TLSs within a similar context. Isolating immune cells specifically from TLSs is not feasible; however, histological analysis reveals that the majority of lymphocytes in our mice reside within TLSs. Thus, we used single-cell RNA sequencing (scRNA-seq) of CD45^+^ cells isolated from livers of 6-month-old DEN-treated IKK RORc Het (control) and IKK RORc KO mice followed by immunohistochemical validation of key findings. Immune cell populations from both genotypes included B cells, plasma cells, CD4 and CD8 T cells, NK cells, monocytes/macrophages, and granulocytes (Fig. S11A).

We observed a significant enrichment of CD8 T cells in IKK RORc KO mice (Fig. 4A-C). Cluster 0 encompassing CD8 effector (CD8_EFF_) cells with both exhaustion (PD1, Lag3, Tox, and Ikzf2) and effector (Perforin, Granzyme A, B, K and IFNg) markers, was the most enriched (15.1 fold) in IKK RORc KO mice (Fig. 4B,C and Fig. S11B). Other enriched CD8 clusters in IKK RORc KO mice included CD8 resident memory (CD8_RM_) cells, expressing the anti-tumor cytokines IFNg and TNF, CD8 memory (CD8_MEM_) cells, and proliferating CD8 cells. In contrast, naïve-like and central memory CD8 cells were reduced in IKK RORc KO mice (Fig. 4B,C and Fig. S11B). The reduction in CD8 naïve-like cells and enrichment of effector/memory CD8 cells indicates a pronounced active process of antigen recognition and differentiation in IKK RORc KO mice.

**Figure 4:**
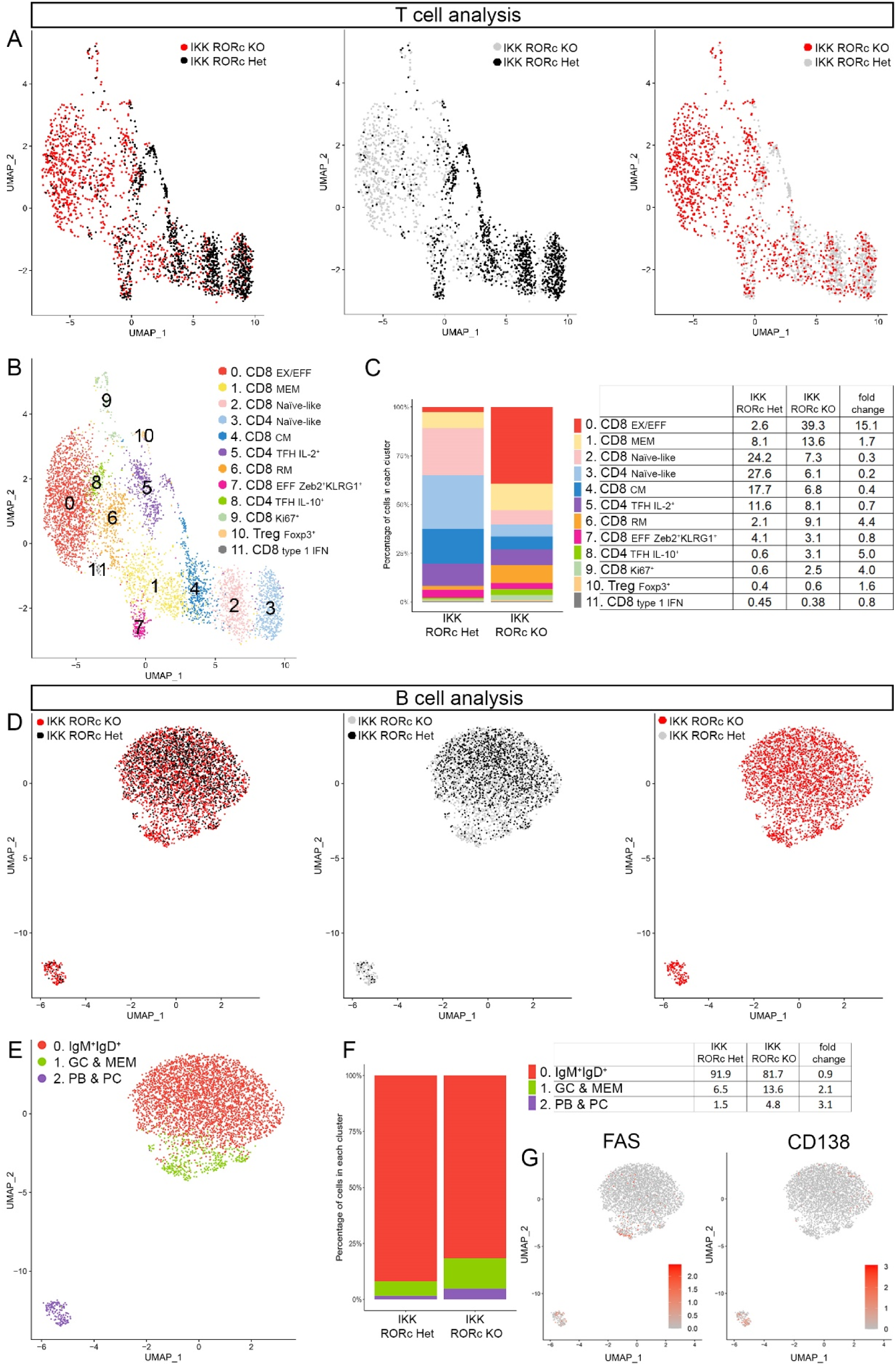
Exhausted yet cytotoxic CD8 cells and class-switched B cells are enriched in IKK RORc KO livers. scRNA-seq analysis of CD45^+^ cells isolated from livers of DEN-injected 6 months old IKK RORc Het mice (pool of 3 mice) and IKK RORc KO mice (2 mice). **(A)** Uniform manifold approximation and projections (UMAPs) restricted to CD3^+^ T cells showing the distribution of cells between IKK RORc Het (black) and IKK RORc KO (red) (left). T cell distribution of IKK RORc Het (black) on top of IKK RORc KO (gray) (middle), and IKK RORc KO (red) on top of IKK RORc Het (gray) (right). **(B)** Same UMAP as A, showing T cells cluster identity. **(C)** Diagram and table showing the distribution of T cells across clusters of each genotype, represented as percentages of total T cells. Fold change for selected clusters= IKK RORc KO/IKK RORc Het. **(D)** UMAP restricted to B cells showing the distribution of cells between IKK RORc Het (black) and IKK RORc KO (red) (left). B cell distribution of IKK RORc Het (black) on top of IKK RORc KO (gray) (middle), and IKK RORc KO (red) on top of IKK RORc Het (gray) (right). **(E)** Same UMAP as D, showing B cells cluster identity. **(F)** Diagram and table showing the distribution of B cells across clusters of each genotype, represented as percentages. Fold change for selected clusters= IKK RORc KO/IKK RORc Het. **(G)** UMAP restricted to B cells showing normalized UMI counts of Fas (GC B cells marker) and CD138 (plasma cells marker) (high expression levels – red, low expression levels – gray). In A,B,D,E the number of cells in IKK RORc KO was randomly down-sampled, in order to show equal numbers of cells from IKK RORc Het and IKK RORc KO. Abbreviations: CD8_EX/EFF_-CD8 exhausted/effector, CD8_EFF_-CD8 effector, CD8_MEM_-CD8 memory, CD8_CM_-CD8 central memory, CD4_TFH_-CD4 T follicular helper, CD8_RM_-CD8 resident memory, T_REG_-T regulatory, GC-germinal center, MEM-memory, PB-plasmablast, PC-plasma cells.

Examining CD4 cells, we found a decrease in naïve-like cells in IKK RORc KO mice compared to controls. Two distinct CD4 clusters were identified: a pro-inflammatory cluster (IL-7r^+^ IL-2^+^) and an anti-inflammatory cluster (IL-7r^-^ IL-10^+^) expressing markers of T follicular helper (T_FH_) cells, the latter being enriched in IKK RORc KO mice (Fig. 4B,C and Fig. S11B). Regulatory T cells (Tregs) expressing Foxp3 did not change significantly between the genotypes (Fig. 4B,C and Fig. S11B). Interestingly, very few RORc^+^ cells were identified in a distinct part of cluster 5 in control mice (Fig. S11B) that seems to be CD3^+^CD4^-^CD8^-^, which suggests that part of the RORcECs possibly represent γδT cells, although we cannot rule out the presence of ILC3 (CD3^-^ CD4^+^) or Th17 (CD3^+^ CD4^+^) cells.

The main populations in the B cell lineages were: unswitched IgM^+^IgD^+^ B cells (Fcer2a^+^ naïve-like cells), germinal center (GC) B cells (Fas^+^, AID^+^, CD86^+^, CD83^+^, Cxcr4^+^, Myc^+^, Cyclin-D2^+^, Bcl2A1b^+^; (Holmes et al., 2020)), memory B cells (CD80^+^, CD73^+^, Cxcr3^+^, ApoE^+^, Scimp^+^, Zbtb20^+^, IgG1^+^, IgG2b^+^; (Yewdell et al., 2021)), plasmablasts (Blimp1^+^ and IRF4^+^) and plasma cells (CD19^-^, CD138^+^, IgA^+^, Jchain^+^, IgG2b^+^, BCMA^+^, Xbp1^+^; (Holmes et al., 2020)) (Fig. 4D-G and Fig. S12). The GC B cell/memory B cell cluster was enriched in IKK RORc KO mice, suggesting increased memory B cell development. The plasmablast/plasma cell cluster was also enriched in IKK RORc KO mice, indicating heightened differentiation into IgA- and IgG-secreting plasma cells (Fig. 4D-G and Fig. S12).

Overall, IKK RORc KO livers exhibited increased populations of CD8 cells with an exhausted phenotype that retained cytotoxic potential, CD4 cells with regulatory functions, and GC B cells that contribute to memory B cell and plasma cell generation. These findings imply an ongoing process of antigen recognition and potential targeting of cancer antigens within the IKK RORc KO background, potentially explaining the anti-tumorigenic phenotype of TLSs in this context.

### RORc expressing cells control the abundance and phenotype of CD8 effector and memory T cells

To further examine T cell states, also at the protein level, we employed cytometry by time-of-flight (CyTOF) analysis. Consistent with the scRNA-seq results, 6 months old DEN-injected IKK RORc KO mice exhibited increased abundance of CD8 cells compared to IKK RORc Het mice. Specifically, CD8_EFF_, CD8_MEM_, and CD8_RM_ populations were enriched in IKK RORc KO mice (Fig. 5A,B and Fig. S13A-C). Conversely, CD4 memory (CD4_MEM_), Tregs, and T_H2_ cells decreased in IKK RORc KO mice, while T_H1_, T_FH_ and tissue-resident memory (CD4_RM_) cells increased (Fig. 5C,D and Fig. S13A). The population of CD3^+^CD8^-^CD4^-^ [CD3 double negative (CD3_DN_)] cells, potentially including RORc^+^ γδT cells, also decreased in IKK RORc KO mice (Fig. 5A,E). Additionally, a population of plasmablasts (CD19^-/+^ IRF4^+^ Ki67^+^) was enriched in IKK RORc KO mice (Fig. 5F,G), indicating ongoing antigen recognition in TLSs.

**Figure 5:**
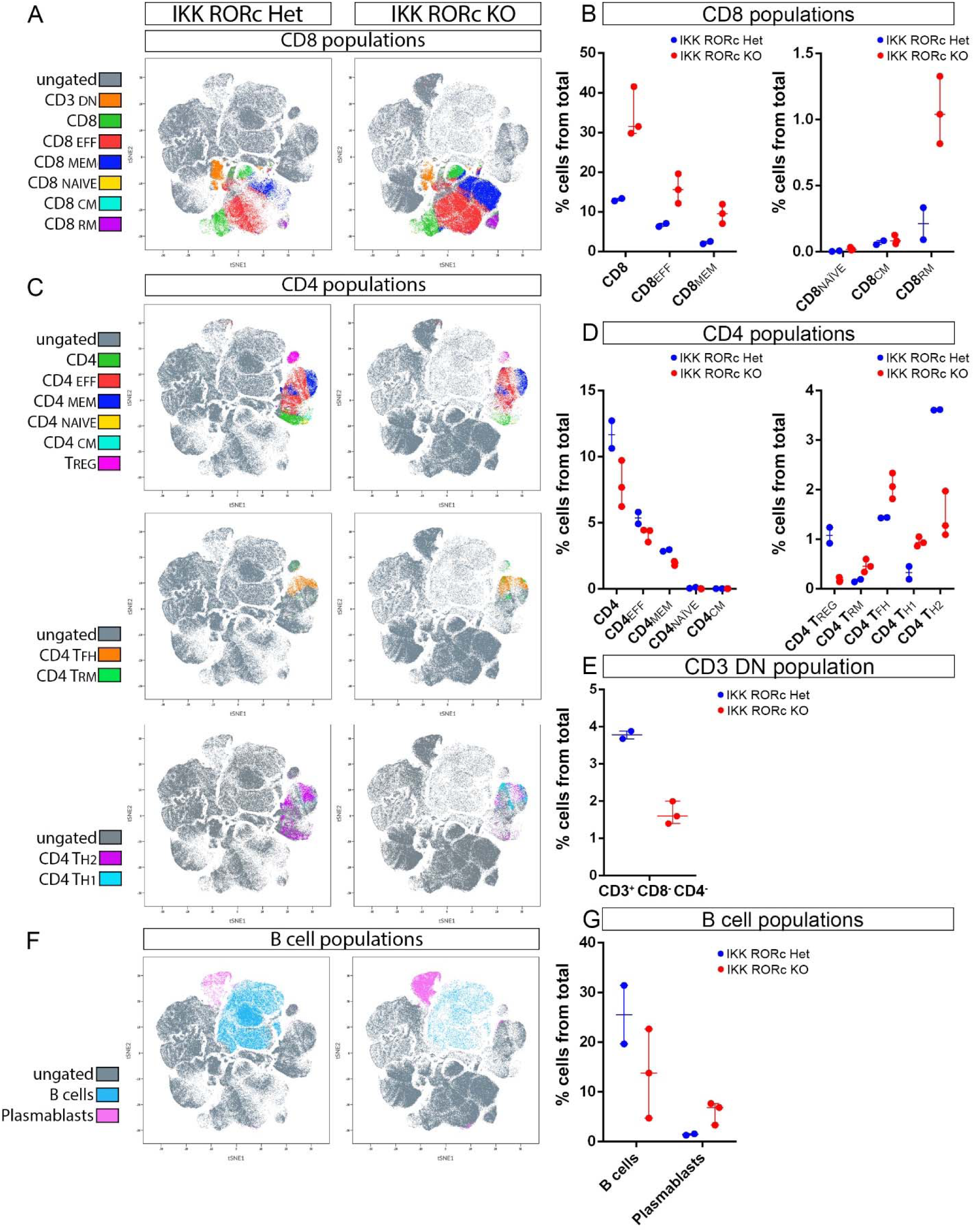
Enrichment of effector and memory CD8 cells and shift from T_H2_ to T_H1_ phenotype of CD4 cells in IKK RORc KO livers. Immune cells were isolated from livers of DEN-injected 6 months old control mice (n=2, each representing a pool of 3 IKK RORc Het or 4 IKK mice) and IKK RORc KO mice (n=3, each from a single mouse) and analyzed using CyTOF. **(A)** tSNE plots highlighting CD8 cells. **(B)** Percentages of CD8 cell subsets from the total immune cell population in IKK RORc Het compared to IKK RORc KO. CD8_EFF_ (CD44^+^ CD262L^-^ CCR7^-^ IL7Ra^-^), CD8_MEM_ (CD44^+^ CD262L^-^CCR7^-^ IL7Ra^+^) and CD8_RM_ (CD103^+^CD69^+^) are increased in IKK RORc KO. **(C)** tSNE plots highlighting CD4 populations. **(D)** Percentages of CD4 cell subsets from the total immune cell population in IKK RORc Het compared to IKK RORc KO. CD4_RM_ (CD103^+^CD69^+^), CD4 T_FH_ (ICOS^+^) and CD4 T_H1_ (Tbet^+^) are increased, while Tregs (Foxp3^+^) and CD4 T_H2_ (Gata3^+^) are decreased in IKK RORc KO. **(E)** Percentages of CD3^+^ CD8^-^ CD4^-^ double negative (DN) cells are decreased in IKK RORc KO (CD3_DN_ cluster is shown in the tSNE in panel A). **(F,G)** B cells (CD19^+^) and plasmablasts (CD19^-/+^ IRF4^+^ Ki67^+^) populations and their percentages. Abbreviations: CD3_DN_-CD3 double negative, CD8_NAÏVE_-CD8 naïve-like, CD4_NAÏVE_-CD4 naïve-like, CD4_EFF_-CD4 effector, CD4_MEM_-CD4 memory, CD4_CM_-CD4 central memory, CD4_RM_-CD4 resident memory, T_H1_-T helper 1, T_H2_-T helper 2. Other abbreviations as in Fig. 4 legend.

Two types of exhausted T cells have been identified in the tumor microenvironment: progenitor cells (PD1^+^Tbet^+^Eomes^-^) with greater polyfunctionality and persistence, and terminally exhausted cells (PD1^+^Tbet^-^Eomes^+^) with enhanced cytotoxicity but reduced survival. The progenitor population gives rise to terminally exhausted T cells, mediates enhanced tumor control and retains responsiveness to PD1 blockade (Miller et al., 2019; Paley et al., 2012; Pauken & Wherry, 2015). We examined PD1 expression in CD8_EFF_ and CD8_MEM_ cells and found that the majority of cells in both populations were PD1 negative (Fig. S14A,B), suggesting they were not exhausted. However, the percent of CD8_MEM_ cells expressing intermediate levels of PD1 (PD1^int^) was lower in IKK RORc KO compared to controls (median=20.5% in controls and 11.1% in IKK RORc KO) (Fig. S14B), suggesting that CD8_MEM_ are less exhausted in IKK RORc KO mice. The proportion of Tbet^+^ Eomes^-^ cells decreases as PD1 levels increase, at least in IKK RORc KO (Fig. S14C). Moreover, the proportion of Tbet^+^ Eomes^-^ PD1^int^ and Tbet^+^ Eomes^-^ PD1^-^ cells was higher in IKK RORc KO compared to control mice in both CD8_EFF_ and CD8_MEM_ cells (Fig. S14A,C,D,H), suggesting that CD8_EFF_ and CD8_MEM_ are less exhausted in the absence of RORcECs.

TCF1 expression has been associated with a progenitor subset of CD8 cells that exhibit better tumor growth control (Kurtulus et al., 2019; Sade-Feldman et al., 2018; Siddiqui et al., 2019). In both IKK RORc KO and control mice, the majority of PD1^high^, PD1^int^ and PD1^-^ cells in CD8_EFF_ and CD8_MEM_ populations expressed TCF1, (Fig. S14F,G), supporting the notion that these cells are not terminally exhausted and may contribute to tumor control.

Additionally, we observed higher levels of pS6 (a downstream effector of the mTOR pathway, required for T cell activation and effector functions), and the cytotoxic marker Granzyme B (GzmB), in exhausted PD1^high^ CD8 cells from IKK RORc KO mice, indicating retained effector functions (Fig. S14C,D,H). Furthermore, the percentage of Gata3-expressing CD8_MEM_ cells (Gata-3 being a driver of T cell dysfunction (Singer et al., 2016)) was lower in IKK RORc KO mice (Fig. S14E,H), suggesting improved functionality compared to controls.

In summary, RORcECs negatively regulate CD8 cell abundance and favor a more exhausted and less cytotoxic CD8 phenotype. RORcECs also promote a T_H2_ over T_H1_ phenotype of CD4 cells and negatively regulate plasmablasts. These changes in the immune phenotype of hepatic lymphocyte populations in IKK RORc KO mice may contribute to an anti-tumorigenic environment.

### Anti-tumorigenic TLSs are enriched in exhausted, yet proliferative, CD8 cells with a progenitor-like phenotype

Both scRNA-seq and CyTOF analyses revealed an increased presence of CD8 cells in the livers of IKK RORc KO mice compared to control mice. Immunohistochemistry staining of liver sections confirmed this enrichment within TLSs. CD8 cell abundance was significantly 3-fold higher in TLSs of IKK RORc KO mice (Fig. 6A,B), as well as in the liver parenchyma (data not shown). Additionally, the exhaustion state of these CD8 cells was assessed by co-staining for CD8, PD1, and the proliferation marker Ki67. We observed a significant increase in CD8^+^ PD1^+^ (6.5-fold), CD8^+^ Ki67^+^(4.8-fold), and CD8^+^ PD1^+^ Ki67^+^ (5-fold) cells in TLSs of IKK RORc KO mice compared to controls (Fig. 6A,B). Thus, TLSs in IKK RORc KO mice harbor a greater number of exhausted, yet proliferative, CD8 cells. The abundance of CD4 cells in TLSs was unchanged (Fig. S15A,C). Moreover, the majority of CD8 cells expressed the progenitor marker TCF1 (52.7% in controls and 70% in IKK RORc KO), with a 4-fold and 5-fold increase in CD8^+^ TCF1^+^ and CD8^+^ PD1^+^ TCF1^+^ cells, respectively, in TLSs of IKK RORc KO mice (Fig. 6A,C), which reflects the total increase in CD8 cells.

**Figure 6:**
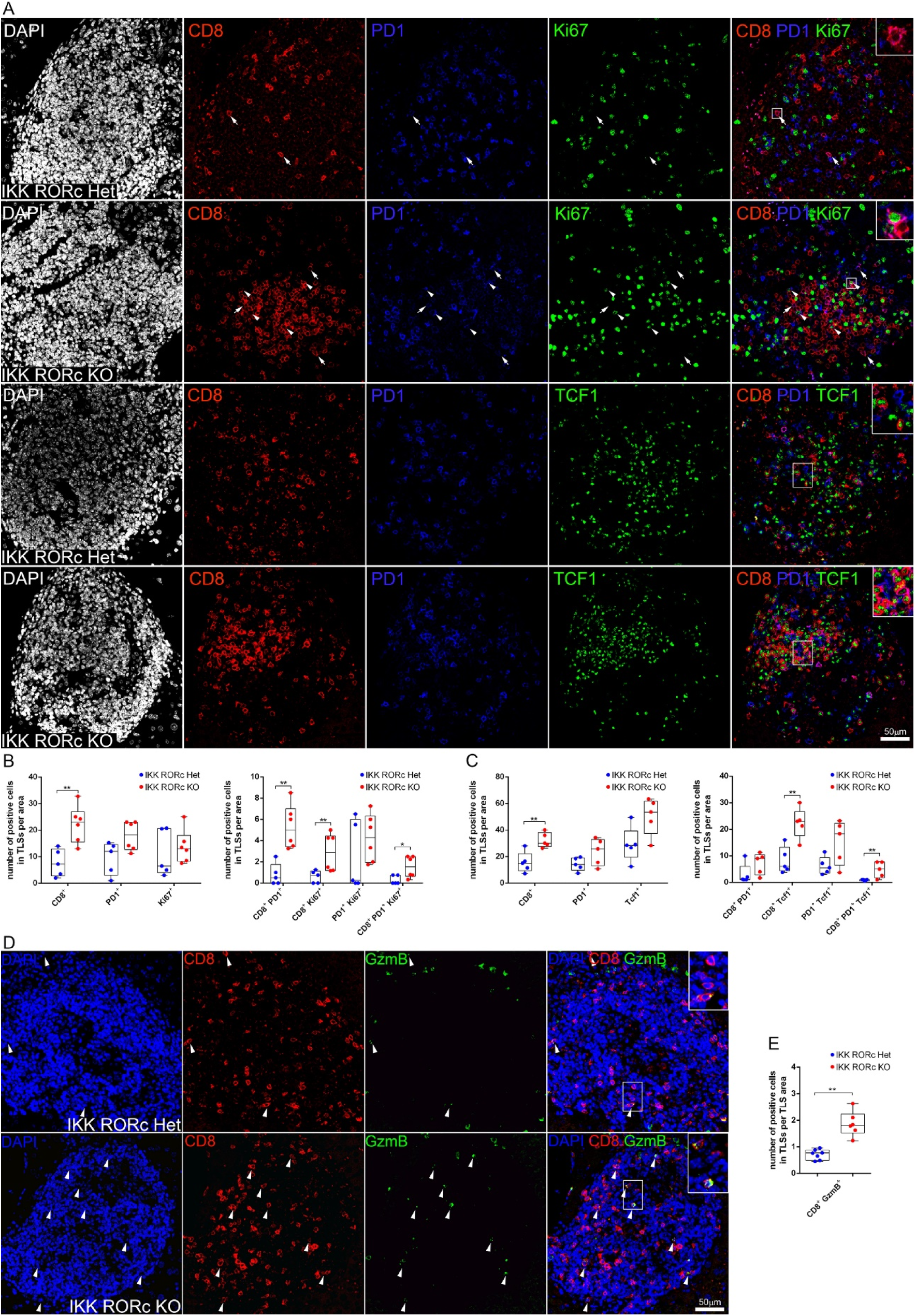
Proliferating and progenitor-like CD8 cells expressing PD1 are enriched within TLSs of IKK RORc KO. **(A)** Triple immunofluorescent staining for the indicated markers in TLSs of IKK RORc Het and IKK RORc KO mice. In the two upper panels, arrows point to CD8^+^ PD1^+^ cells and arrowheads point to CD8^+^ PD1^+^ Ki67^+^ cells. Cells co-expressing CD8 (red) and PD1 (blue) are seen in pink. Insets correspond to areas in white boxes. **(B)** Number of cells expressing CD8, PD1 or Ki67 (left) and their combinations (right) within TLSs. **(C)** Number of cells expressing CD8, PD1 or TCF1 (left) and their combinations (right) in TLSs. **(D)** Immunofluorescent staining for the indicated markers in TLSs of IKK RORc Het and IKK RORc KO mice. CD8^+^ GzmB^+^ cells are marked by arrowheads. **(E)** Number of cells expressing CD8^+^ GzmB^+^ in TLSs. All analyses were done in DEN-injected 6 months old mice. Statistical test applied: Mann-Whitney.

This suggests that within TLSs, the abundance of CD8 cells that are not terminally exhausted and have a more progenitor-like phenotype is higher in IKK RORc KO. Furthermore, an increase in CD8^+^GzmB^+^ cytotoxic cells (Fig. 6D,E) and CD8^+^CD103^+^ CD8_RM_ cells (Fig. S13B,C) was observed within TLSs of IKK RORc KO mice, reflecting the total increase in CD8 cells. There was no difference in the numbers of either Foxp3^+^ or Foxp3^+^ PD1^+^ Treg cells in TLSs of controls compared to IKK RORc KO mice (Fig. S15B,C). The previous decrease in Treg numbers seen in the CyTOF results may reflect changes in non-TLS regions of the liver. Very few myeloid-derived suppressor cells (MDSCs) were detected in TLSs and no difference was seen between IKK RORc KO and control mice (Fig. S15D).

### Anti-tumorigenic TLSs show an augmented B cell response

According to the scRNA-seq and CyTOF analyses, there was no significant quantitative difference in the percentage of B cells. However, both methods revealed an increase in plasmablasts (precursors of plasma cells) in IKK RORc KO mice compared to controls (Fig. 4D-G and Fig. 5F,G). GC B cells and plasma cells were also enriched in IKK RORc KO mice according to the scRNA-seq data (Fig. 4E-G), indicating an augmented B cell response in livers of IKK RORc KO mice. Further validation was conducted by characterizing the TLS maturation state using markers of GC B cells. Proliferating GC B cells, identified by peanut agglutinin (PNA) binding (CD19^+^PNA^+^Ki67^+^), were observed in both control and IKK RORc KO mice within TLSs. However, the percentage of GC-positive TLSs was higher in IKK RORc KO mice (Fig. 7A-C), suggesting a greater maturity of TLSs in IKK RORc KO mice. Another GC B cell marker is activation induced cytidine deaminase (AID), which is required for isotype switching and affinity maturation. TLSs harboring AID-expressing cell clusters were seen in both controls and IKK RORc KO mice but their percent was higher in IKK RORc KO mice (Fig. 7D-F). Thus, TLSs of IKK RORc KO mice have an enhanced capacity to generate GCs and reach higher maturity compared to their control counterparts. These results suggest that more plasma cells and antibodies may be generated in IKK RORc KO mice. Plasma cells, identified by CD138 staining, were localized at different densities, mainly surrounding and adjacent to TLSs and in some cases within TLSs of all sizes, with very few plasma cells in the parenchyma (Fig. 7G,H). Qualitative assessment indicated a higher percentage of TLSs with medium and high densities of plasma cells in IKK RORc KO mice (Fig. 7I). Furthermore, staining for IgA and IgG antibodies co-localized with CD138 staining and demonstrated a quantitative increase in IKK RORc KO mice compared to controls (Fig. 7J-M), supporting the notion that plasma cells producing IgA and IgG antibodies are more abundant in IKK RORc KO mice, and are likely generated by more mature GC-harboring TLSs.

**Figure 7:**
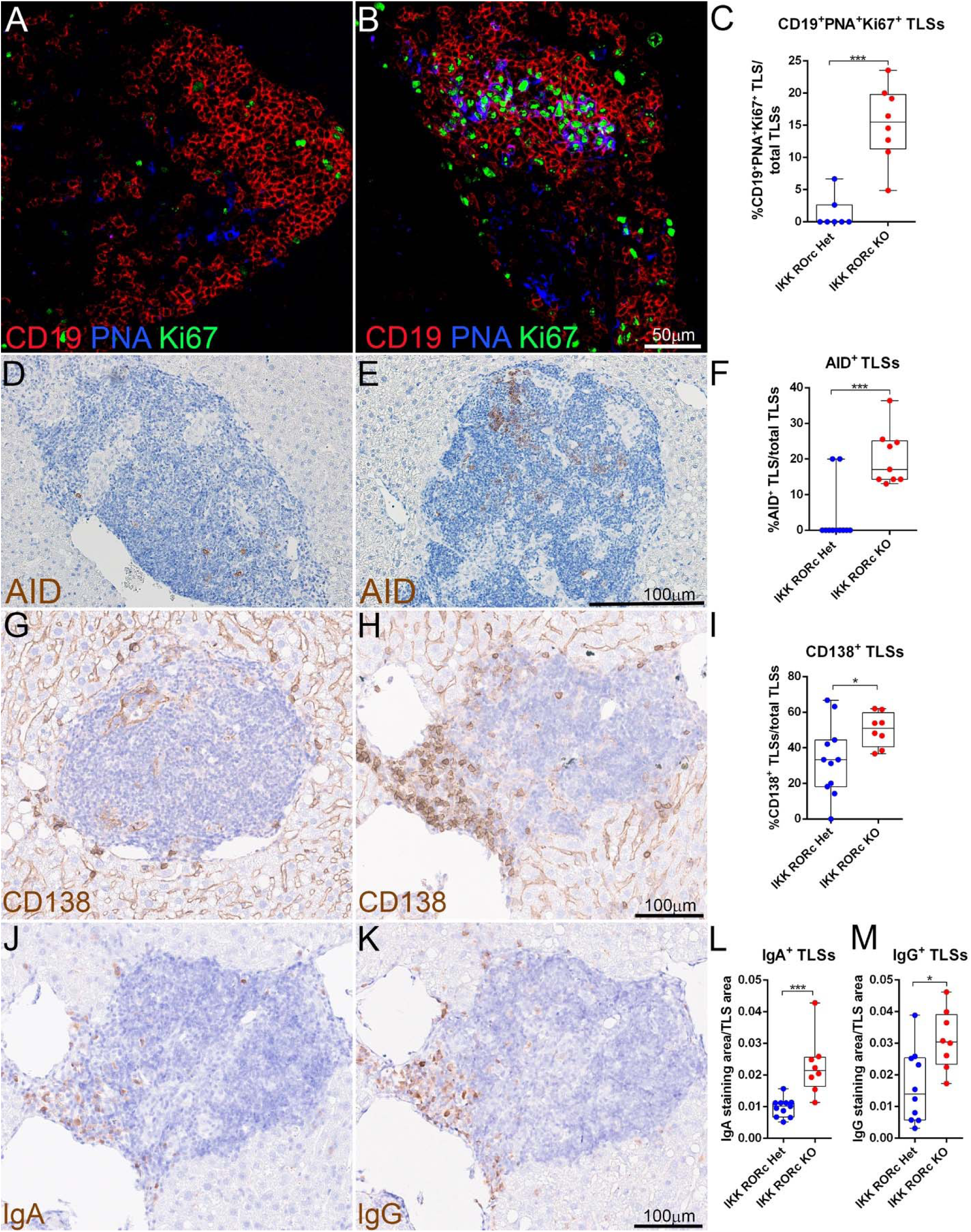
IKK RORc KO mice show increased B cell response. Examples of GC-negative **(A)** or - positive **(B)** TLSs, assessed by the presence of cell clusters triply positive for CD19, PNA and Ki67. **(C)** Quantification of the percent of TLSs containing CD19^+^ PNA^+^ Ki67^+^ cell clusters, in IKK RORc Het and IKK RORc KO mice. Examples of GC-negative **(D)** or -positive **(E)** TLSs, assessed by the presence of cell clusters expressing AID. **(F)** Quantification of the percent of TLSs containing AID^+^ cell clusters in IKK RORc Het and IKK RORc KO mice. Examples of low density **(G)** or high density **(H)** plasma cell distribution in TLSs, assessed by staining for CD138 (medium plasma cell density is not shown); CD138 is also expressed in a sinusoidal pattern along with immunostaining of hepatocytes and biliary duct epithelium, interfering with its quantitation. **(I)** Qualitative quantification of the percent of TLSs containing high and medium density of CD138^+^ plasma cells. Serial sections of tissue seen in panel (H), stained for IgA **(J)** or IgG **(K)** and the corresponding quantification of staining area relative to TLS area (**L,M**, respectively). All analyses were done in DEN-injected 6 months old mice. Statistical test applied: Mann-Whitney.

### B cells protect against iCCA formation

IKK RORc KO mice exhibit enrichment of CD8 cells, altered subsets of CD4 cells, and increased B cell response in TLSs, all of which could account for the reduced iCCA formation in this background. We therefore evaluated the role of immune cell populations in tumor formation, by cell depletion experiments in 6 months old DEN-injected IKK RORc Het (control) and IKK RORc KO mice (See Fig. 2 for evaluation of TLS development following the same immune cell depletion regimes). In control mice, depletion of CD4, CD8 or B cells did not affect HCC abundance compared to isotype treated mice (Fig. 8A-D, Fig. S16A-B’,F-H, Fig. S17A-B’, F-H and Fig. S18A-B’, F-H). Depletion of CD4 or CD8 cells did not affect iCCA tumors (Fig. 8E,F, Fig. S16A-E and Fig. S17A-E). However, the iCCA/TLS ratio was significantly increased following CD4 (but not CD8) depletion (Fig. 8H,I), suggesting CD4 cells contribute to the formation of anti-tumorigenic TLSs. Importantly, depletion of B cells significantly increased iCCA numbers and the iCCA/TLS ratio (Fig. 8G,J and Fig. S18A-E), suggesting B cells limit TLS-related iCCA development. Smaller iCCAs showed a more pronounced increase compared to larger ones, suggesting that B cells inhibit iCCA formation rather than growth (Fig. S18E). B cells may exert their anti-tumor effects through antigen presentation to CD8 cells or by producing anti-tumor antibodies; yet the lack of impact on iCCA formation after depleting CD8 cells makes the former possibility less likely. We thus tested if B cell depletion reduced antibody generation. Primary and secondary antibody responses are abrogated following CD20 depletion, while long-lived plasma cells are unaffected (DiLillo et al., 2008). Our prolonged two months CD20 depletion regime significantly reduced the abundance of CD138^+^ plasma cells (Fig. 8K and Fig. S19A). This effect was more pronounced for IgG (and less so IgA) positive plasma cells (Fig. 8L,M and Fig. 19B,C). These results suggest a diminished capacity to generate new antibody-secreting cells in TLSs.

**Figure 8:**
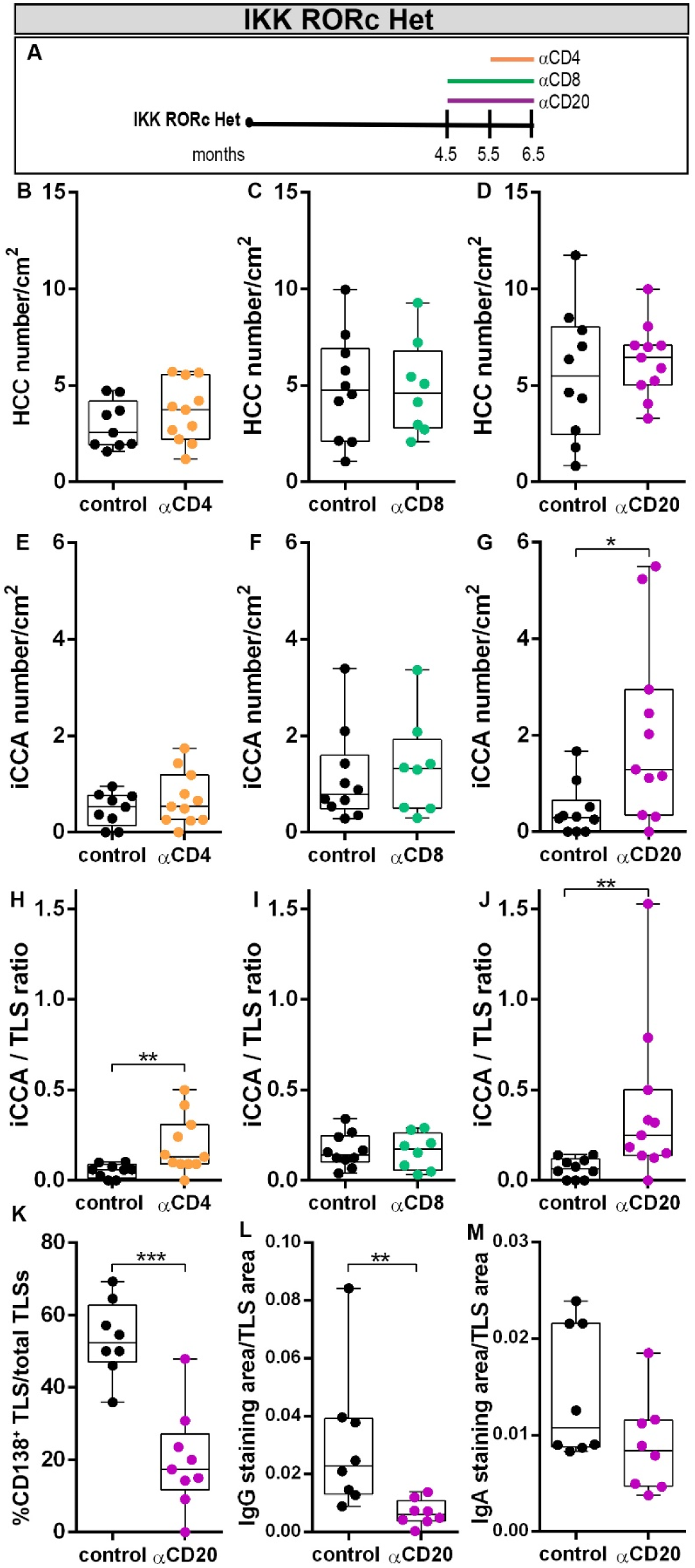
B cells protect against iCCA formation. Timeline of cell depletion in DEN-injected IKK RORc Het mice **(A)**. HCC number in CD4- **(B)**, CD8- **(C)** and CD20-depleted **(D)** IKK RORc Het mice. iCCA number in CD4- **(E)**, CD8- **(F)** and CD20-depleted **(G)** IKK RORc Het mice. Ratio of iCCA numbers to TLS numbers in CD4- **(H)**, CD8- **(I)** and CD20- **(J)** depleted IKK RORc Het mice. **(K)** Qualitative quantification of the percent of TLSs containing high and medium density of CD138^+^ plasma cells in CD20 depleted IKK RORc Het mice. Quantification of IgG **(L)** and IgA **(M)** staining area relative to TLS area in CD20 depleted IKK RORc Het mice. Each dot represents an average of all TLSs in a mouse (L,M). Statistical test applied: Mann-Whitney.

Next, we performed the same depletion regimes in IKK RORc KO mice. Of note, IgG2a isotype antibodies, used as controls, increased iCCA numbers in IKK RORc KO mice, hindering their comparison to untreated IKK RORc KO mice. CD8 depletion in IKK RORc KO mice reduced HCC numbers and did not affect iCCA or the iCCA/TLS ratio (Fig. S20A,B,D,F and Fig. S21), suggesting that enriched CD8 cells in this background promote HCC development and do not contribute to TLS-related iCCA development. Depletion of B cells in IKK RORc KO mice did not affect HCC or iCCA numbers (Fig. S20A,C,E and Fig. S22) but increased the iCCA/TLS ratio (Fig. S20G), suggesting that B cells limit TLS-related iCCA formation. Notably, while B cell depletion in the spleen of IKK RORc KO mice was efficient (Fig. S3J), depletion in the liver was less efficient than in control mice (Fig. S4C,D,F,G). Moreover, Plasma cells, IgA- and IgG-expressing cells in TLSs were increased rather than decreased (Fig.20H,I and Fig. S23). These effects probably impede a full evaluation of the contribution of B cells to iCCA formation in IKK RORc KO mice, yet are in line with the notion that antibodies may be responsible for curtailing iCCA development.

Altogether, we find that B cells rather than CD8 cells limit TLS-related iCCA development, and that while RORcECs regulate both populations, their ability to promote iCCA development is likely mediated by downregulation of the B cell response.

## Discussion

TLSs have prognostic and predictive value in cancer, mostly predicting a favorable outcome, but are associated with a poor prognosis in several cancers, including liver cancer (Figenschau et al., 2015; Finkin et al., 2015; Joshi et al., 2015; Sautès-Fridman et al., 2019). TLSs contribute to the response to immune checkpoint blockers (ICB) or reflect the tumor microenvironment’s permissiveness to ICBs (Schumacher & Thommen, 2022). Manipulating TLS neogenesis has significant therapeutic implications. Our study demonstrates a genetic manipulation that enhances TLS neogenesis in the context of liver cancer. Mice lacking RORc develop excess TLSs that depend on B cells and CD4 cells for their neogenesis and maintenance, respectively. This manipulation not only increases TLS numbers, but also shifts their phenotype from pro- to anti-tumorigenic. RORcECs inhibit CD8 cell accumulation with an exhausted/cytotoxic phenotype, promote a T_H2_ over T _H1_ phenotype of CD4 cells and inhibit B cell maturation in TLSs. Notably, CD8 cells do not contribute to iCCA control, but an enhanced B cell response likely inhibits iCCA development. This suggests that B cell maturation in TLSs could act as a potential biomarker to differentiate between pro- and anti-tumorigenic TLSs in peritumor regions of iCCA. Moreover, manipulating RORcECs or subsets of CD4 and B cells may regulate TLS abundance and phenotype.

### Indirect regulation of TLS formation by RORc expressing cells

Our study reveals that RORcECs are not essential for TLS neogenesis and surprisingly, they even negatively regulate TLS formation. Our results suggest that either dysregulated CD4 and/or B cells in the liver could account for excess TLS formation in the absence of RORc, suggesting that RORcECs control CD4 or B cell number or functions in TLSs. Interestingly, B cells are required for TLS neogenesis only in the absence of RORc and not in IKK RORc Het mice, suggesting that RORcECs suppress the ability of B cells to facilitate TLS formation. In the absence of RORc, isotype-switched B cells are increased, however it is less likely that they would have a role in TLS neogenesis since they differentiate only as TLSs mature. Thus, although their numbers are not altered, naïve-like/other B cells probably provide the first signals that promote the assembly of TLSs. Notably, B cell regulation of tumor progenitor development within TLSs is independent of CD8 cells. Importantly, CD4 cells are required for TLS maintenance in the presence or absence of RORcECs. Some CD4 populations, including T_FH_ cells (shown to promote TLS formation in human tumors (Chaurio et al., 2022)), are enriched in the absence of RORc and could account for excess TLS formation. Thus, RORcECs control TLS numbers by negatively regulating CD4 cells required for TLS maintenance and/or by inhibiting the potential of B cells to promote TLS neogenesis.

### Indirect regulation of iCCA formation by RORc expressing cells

We observe changes in the profile of liver immune cells in IKK RORc KO mice, supporting an indirect effect on iCCA formation, although a direct effect of RORcECs cannot be ruled out. Spontaneous immune activation has been reported in RORc KO mice, with elevated cytokine levels and increased frequencies of effector memory CD4 and CD8 cells (Wichner et al., 2013). Similarly, we observe a quantitative increase in exhausted/cytotoxic CD8 cells and IFNg-expressing CD8_RM_ cells in anti-tumorigenic TLSs of IKK RORc KO mice. Depletion experiments in IKK RORc Het and IKK RORc KO mice show that CD8 cells do not affect TLS formation or iCCA development. While the enriched CD8 population in IKK RORc KO mice exhibits a mixed phenotype, expressing markers associated with effector function, exhaustion and tissue residency, these CD8 cells may be unable to prevent tumor growth despite their cytotoxic potential. In nonalcoholic steatohepatitis-induced HCC, CD8^+^PD1^+^ T cells, with a similar exhausted/cytotoxic/residency phenotype, did not cause tumor regression but instead contributed to tissue damage (Pfister et al., 2021; Dudek et al., 2021). In agreement with this, CD8 depletion in IKK RORc KO mice reduced HCC formation. Therefore, the increased CD8 populations in IKK RORc KO mice might be incapable of halting iCCA formation and can even promote HCC formation.

We observe a moderate increase in liver GC B cells, plasmablasts, and plasma cells in IKK RORc KO mice, as similarly observed in spleens of RORc KO mice (Wichner et al., 2016). Several recent studies point to the role of TLS-residing B cells in cancer immunology in general and in the response to ICBs in particular (Cabrita et al., 2020; Griss et al., 2019; Helmink et al., 2020; Petitprez et al., 2020). B cell depletion in IKK RORc Het mice increased iCCA numbers and the iCCA/TLS ratio without affecting TLS formation, suggesting a role for B cells in limiting iCCA development. The decrease in plasma cell numbers after prolonged B cell depletion may suggest the involvement of an anti-tumor antibody response. Indeed, recent studies highlight the contribution of TLS-residing B cells to anti-tumor response through the production of tumor-binding antibodies and involves antibody-dependent cellular cytotoxicity (Ng et al., 2023). Understanding the effect of B cell depletion in IKK RORc KO mice is more complicated since it also reduced TLS numbers which directly affect iCCA formation. Therefore, the effect of B cell depletion on tumor formation cannot be uncoupled from its effect on TLS formation. Moreover, incomplete liver B cell depletion efficiency and unexplained higher levels of plasma cells in TLSs, make it hard to define the role of B cells in regulating iCCA formation in the absence of RORc. Yet, the iCCA/TLS ratio is still higher in B cell depleted IKK RORc KO compared to controls, in agreement with an anti-tumor function of B cells. Also, in CD4-depleted IKK RORc Het mice, the impact of cell depletion on tumor formation cannot be separated from its influence on TLS formation. However, the increased iCCA/TLS ratio suggests a role for CD4 cells in inhibiting iCCA development. Potentially, CD4 T_FH_ cells (increased in IKK RORc KO mice) would support B cell differentiation to plasma cells that limit iCCA formation, explaining the effect of both CD4 and B cells on iCCA formation.

We find that B cells have dual opposing roles: 1) Promoting neogenesis of TLSs which harbor tumor progenitors that give rise to iCCA; 2) Limiting iCCA formation. This contradiction can be explained by the involvement of different B cell populations with either pro- or anti-tumor functions at different times. In our model, naïve-like B cells might promote tumor progenitor development initially, while antibody-secreting cells in mature TLSs might limit tumor growth. RORcECs might inhibit the pro-tumorigenic function of naïve-like B cells in early TLSs, and at later time points restrict the development of anti-tumorigenic isotype-switched B cells.

### TLSs in iCCA development

The role of TLSs in iCCA development remains unclear, but recent findings suggest that their abundance within and around tumors is associated with prognosis. In patients with iCCA, high intratumoral TLS abundance is linked to a favorable prognosis, while high peritumoral TLS abundance indicates a poor prognosis, supporting the concept that TLSs can have pro- or anti-tumorigenic properties. T_FH_ cells were more prevalent in intratumoral TLSs, consistent with our mouse data (Ding et al., 2022). Although our mouse model mainly exhibits peritumoral TLSs, their phenotype and cellular composition change upon RORc deletion, suggesting that TLSs do not necessarily need to be located within tumors in order to act in an anti-tumorigenic manner. Treating iCCA patients with a high peritumoral TLS score with RORc inhibitors might transform these TLSs into anti-tumorigenic ones. Moreover, the presence or absence of RORc-expressing immune cells may serve as a marker for the pro- or anti-tumorigenic nature of human hepatic TLSs. The specific identity of RORcECs regulating iCCA formation is yet to be determined, but our data suggest a potential involvement of γδT cells, since CD3 double negative RORcECs were identified in the liver. Interestingly, γδT cells were recently found to be the only immune cell type negatively associated with survival in human iCCA (Zimmer et al., 2022).

TLS induction is crucial for effective cancer treatment, but controlling their nature is equally important. We show that deletion of RORcECs resulted in induction of excess TLSs with anti-tumorigenic potential. Further research is needed to discover whether there is a link between the cells and signals that induce TLS formation and the ways they shape the TLS immune composition and phenotype. While ICBs have revolutionized the treatment of liver cancer, response rates are still low (Llovet et al., 2021). It is thus interesting to speculate that manipulating TLS states could affect responses to ICBs in liver cancer. Additionally, assessing TLS phenotypes, and particularly the prevalence of RORcECs, could serve as a valuable immunotherapy response biomarker in liver cancer.

## Acknowledgments

We thank Shlomi Finkin, Orit Pappo and Michael Berger for their help and advice; Fatima Mushasha, Nasrallah Nasrallah, Bareket Ben-Amos, Abdelmajeed Nasereddin, Idit Shiff, Hasan Sourikh, Eleonora Medvedev and Mohammad Jumaa for excellent technical assistance; Tali Bdolah-Abram for help with the statistical analyses.

## Methods

| Reagent type | Designation | Source | Identifier | Additional information |
| --- | --- | --- | --- | --- |
| Tissue Digestion | Accumax | Sigma/Merck | A7089 | 1 : 2 |
| Tissue Digestion | Liver digest medium | GIBCO | 17703-034 | used at 100% |
| Enzyme | Deoxyribonuclease I from bovine pancreas (DNase) | Sigma/Merck | DN25 | 0.07% |
| DNA stains | Propidium iodide solution | CliniSciences | 30901000-1 | 1 : 500 |
| DNA stains | Trypan blue | Sigma | T8154 | 1 : 2 |
| DNA stains | Hematoxylyn | Kaltek Srl | 1100 |  |
| Chemical | Percoll | Sigma/Merck | P1644-500ML |  |
| Chemical | Diethylnitrosamine (DEN) | Sigma/Merck | N0258 | 10 mg/kg |
| Chemical | Foxp3 / Transcription Factor Staining Buffer Set | eBioscience | 00-5523-00 |  |
| Inhibitor | Brefeldin A | Biolegend | 420601 |  |
| Serum | Donkey serum | Sigma/Merck | D9663 |  |
| Serum | Rat serum | Sigma/Merck | R9759 | 1 : 100 |
| Serum | Fetal Bovine Serum (FBS) | Biological Industries/Sartorius | 04-007-1A |  |
| Serum | Bovine Serum Albumin (BSA) | Sigma/Merck | A3803 |  |
| Antibody | PB-conjugated-anti-CD45 | Biolegend | 109820 | 1 : 500 |
| Antibody | APC-conjugated-anti-F4/80 | eBioscience | 17-4801-82 | 1 : 50 |
| Antibody | CD16/CD32 Monoclonal Antibody (93) (Fc Blocker) | eBioscience | 14-0161-82 | 1 : 100 |
| Antibody | anti-Sox9 | Millipore | AB5535 | 1 : 1000, Citrate pH 6.0 |
| Antibody | anti-E-cadherin | Transduction Laboratories | 610181 | 1 : 200, Citrate pH 6.0 |
| Antibody | anti-CD44v6 | eBioscience | BMS145 | 1 : 500, Citrate pH 6.0 |
| Antibody | anti-HNF4a | Abcam | ab201460 | 1 : 2000, Tris pH 9.0 |
| Antibody | anti-CK19 | DSHB | TROMA-III CK19 | 1 : 25, Citrate pH 6.0 |
| Antibody | anti- CD8 | eBioscience | 14-0808-80 | 1 : 50 , Citrate pH 6.0 |
| Antibody | anti-CD4 | eBioscience | 14-9766-80 | 1 : 50 , Citrate |
| Antibody | anti-CD19 | eBioscience | 14-0194-82 | 1 :500, Citrate pH 6.0 |
| Antibody | anti-mouse PD1 | R&D systems | AF1021 | 1 : 100, Citrate pH 6.0 |
| Antibody | anti- Ki67 | Abcam | ab15580 | 1 : 200, Citrate pH 6.0 |
| Antibody | anti-Foxp3 | eBioscience | 14-5773 | 1 : 50, EDTA pH 8.0 |
| Antibody | anti-RORc | Abcam | ab207082 | 1 : 1000, Tris pH 9.0 |
| Antibody | anti-Tcf1/7 | Cell Signaling | 2203 | 1 : 100, Citrate pH 6.0 |
| Antibody | anti-Ly6G | BD pharmingen | 551459 | 1 : 250, Citrate |
| Antibody | anti-GzmB | Cell Signaling | 44153 | 1 : 100, Citrate pH 6.0 |
| Antibody | anti-CD103 | Abcam | ab224202 | 1 : 1000 Tris pH 9.0 |
| Antibody | anti-IgA | AbD Serotec | STAR137P | 1 : 50, Citrate pH 6.0 |
| Antibody | anti-IgG | AbD Serotec | STAR13B | 1 : 50, Citrate pH 6.0 |
| Antibody | anti-PNA | Vector | VE-CL-1075-1 | 1 : 1000, Citrate pH 6.0 |
| Antibody | anti-AID | eBioscience | 14-5959-82 | 1 : 50, Citrate pH 6.0 |
| Antibody | HRP-conjugated anti-Rat | Histofine | 11-414311F |  |
| Antibody | HRP-conjugated anti-Rabbit | Vector Laboratories | VE-MP-7401-50 |  |
| Antibody | Cy5-conjugated anti-mouse | Jackson immunoresearch | 711-175-152 | 1 : 500 |
| Antibody | Cy3-conjugated anti-rat | Jackson Immunoresearch | 712-165-153 | 1 : 500 |
| Antibody | Cy5-conjugated anti-Goat | Jackson Immunoresearch | 705-175-147 | 1 : 500 |
| Antibody | Cy2-conjugated anti-Rabbit | Jackson Immunoresearch | 711-255-152 | 1 : 200 |
| Antibody | Murine IgG2a antibody GP120:9674 | Genentech |  | 10mg/kg |
| Antibody | Anti-mouse CD20 mAb [clone 5D2], isotype IgG2a | Genentech |  | 10mg/kg |
| Antibody | anti-CD20 [18B12], isotype IgG2a | Absolute Antibody | Ab01110-2.0 | 10mg/kg |
| Antibody | Anti-Fluorescein [4-4-20 (enhanced)] IgG2a Kappa | Absolute Antibody | Ab00102-2.0 | 10mg/kg |
| Antibody | Anti-CD8 alpha/Lyt-2 [YTS 169.4], isotype IgG2a | Absolute Antibody | Ab00166-2.0 | 10mg/kg |
| Antibody | InVivoPlus anti-mouse CD4 [clone GK 1.5], isotype IgG2b | BioXcell | BP0003-1 | 10mg/kg |
| Antibody | InVivoPlus rat IgG2b, anti-keyhole limpet hemocyanin | BioXcell | BP0090 | 10mg/kg |
| Antibody | Pacific Blue anti-mouse CD8a [clone 53-6.7] | Biolegend | 100725 | 1 : 100 |
| Antibody | Alexa Fluor 488 anti-mouse CD4 [clone RM4-5] | Biolegend | 100532 | 1 : 800 |
| Antibody | APC anti-mouse CD3 [clone 17A2] | Biolegend | 100235 | 1 : 100 |
| Antibody | PE anti mouse CD19 [clone 6D5] | Biolegend | 115507 | 1 : 100 |
| Antibody | 111Cd-CD44 | Biolegend | BLG-103002 |  |
| Antibody | 143Nd-CD69 | FLUIDIGM | 3143004B |  |
| Antibody | 145Nd-CD4 | FLUIDIGM | 3145002B |  |
| Antibody | 148Nd-CD103 | Biolegend | BLG-121402 |  |
| Antibody | 149Sm-CD19 | FLUIDIGM | 3149002B |  |
| Antibody | 152Sm-CD3e | FLUIDIGM | 3152004B |  |
| Antibody | 153Eu-CD8a | FLUIDIGM | 3153012B |  |
| Antibody | 159Tb-PD1 | FLUIDIGM | 3159024B |  |
| Antibody | 160Gd-CD62L | FLUIDIGM | 3160008B |  |
| Antibody | 164Dy-CCR7 | FLUIDIGM | 3164013A |  |
| Antibody | 175Lu-IL7Ra | FLUIDIGM | 3175006B |  |
| Antibody | 176Yb-ICOS | FLUIDIGM | 3176014B |  |
| Antibody | 113Cd-Ki67 | Biolegend | BLG-652402 |  |
| Antibody | 115In-Eomes | eBioscience | 14-4875-82 |  |
| Antibody | 155Gd-IRF4 | FLUIDIGM | 3155014B |  |
| Antibody | 158Gd-Foxp3 | FLUIDIGM | 3158003A |  |
| Antibody | 161Dy-Tbet | FLUIDIGM | 3161014B |  |
| Antibody | 167Er-Gata3 | FLUIDIGM | 3167007A |  |
| Antibody | 171Yb-GranzymeB | FLUIDIGM | 3171002B |  |
| Antibody | 172Yb-pS6 | FLUIDIGM | 3172008A |  |
| Antibody | 173Yb-TCF1 | R&D systems | MAB8224 |  |
| Other | FDM barcode 2,8,15,18,19,20 | FLUIDIGM | 201060 |  |
| Other | gentleMACS™ Dissociator | Miltenyi Biotec | 130-093-235 |  |
| Other | Wash buffer | Zytomed Systems | ZUC020-50 |  |
| Other | Protein block | Abcam | ab64226 |  |
| Other | ImmPACT DAB Kit | Vector Laboratories | VE-SK-4105 |  |
| Other | Maxpar PBS | FLUIDIGM | 201058 |  |
| Other | Maxpar MCP9 Antibody Labeling Kit, 111Cd | FLUIDIGM | 201111A |  |
| Other | Maxpar MCP9 Antibody Labeling Kit, 113Cd | FLUIDIGM | 201113A |  |
| Other | MIBItag Conjugation Kit_115In | IONpath | 600115 |  |
| Other | MIBItag Conjugation Kit_173Yb | IONpath | 600173 |  |
| Other | Maxpar Cell Staining Buffer | FLUIDIGM | 201068 |  |
| Other | Cell-ID 20-Plex Pd Barcoding Kit | FLUIDIGM | 201060 |  |
| Other | Cell-ID Cisplatin | FLUIDIGM | 201064 | 1 : 4000 |
| Other | Cell-ID Intercalator-Ir—125 μM | FLUIDIGM | 201192A | 1 : 1000 |
| Other | Maxpar Water | FLUIDIGM | 201069 |  |
| Other | EQ Four Element Calibration Beads | FLUIDIGM | 201078 |  |

### Mice

All animal studies were approved by the Hebrew University Institutional Animal Care and Use Committee. Previously described ROSA26-LSL-IKKβ(EE) mice (Sasaki et al., 2006) were bred with Albumin (Alb)-Cre mice obtained from Jackson Laboratory (Bar Harbor, Maine, stock # 003574) to generate Alb-IKKβ(EE)^Hep^ mice as described previously (Finkin et al., 2015). RORc knock out mice (B6.129P2(Cg)-*Rorc^tm2Litt^*/J) and floxed RORc mutant (B6(Cg)-*Rorc^tm3Litt^*/J) mice were purchased from Jackson Laboratory (stock # 007572 and 008771, respectively), and bred into Alb-IKKβ(EE)^Hep^ mice. The IKK RORc KO colony was kept by mating IKK RORc KO males to IKK RORc Het females. Genotyping was done according to the suppliers’ protocols:

Alb-Cre: https://www.jax.org/Protocol?stockNumber=003574&protocolID=26917

IKKβ(EE): https://www.jax.org/Protocol?stockNumber=008242&protocolID=27052

Rorc^tm2Litt^: https://www.jax.org/Protocol?stockNumber=007572&protocolID=23307

Rorc^tm3Litt^: https://www.jax.org/Protocol?stockNumber=008771&protocolID=27794

All mice were of a pure C57BL/6J genetic background and were bred and maintained in specific pathogen-free conditions. Only male mice were used for experiments. Animals were sacrificed by a lethal dose of anesthesia and perfused through the left ventricle with heparinized PBS. For hepatocarcinogenesis, mice were injected intraperitoneally (i.p) with 10 mg/kg Diethylnitrosamine (DEN) at 14 days of age. Mice were observed for development of tumors at 6.5 months of age.

### *In Vivo* cell depletion

Littermates were treated with depleting antibodies and matching isotype controls. Cell depletion experiments were compared only to isotype treated controls since TLS numbers were increased around 1.5 fold by isotype controls alone. IgG2a treatment alone also caused an increase in iCCA numbers compared to untreated mice, hindering their comparison. All antibodies were used at a concentration of 10mg/kg. Mice were injected i.p twice a week with anti-CD4 antibodies (clone GK 1.5, BioXcell), once a week with anti-CD8 antibodies (clone YTS 169.4, Absolute Antibody) and once every two weeks with anti-CD20 antibodies (clone GP120:9674, Genentech or 18B12, Absolute Antibody). Mice were sacrificed 3 to 7 days after the last injection. IgG2b (anti-keyhole limpet hemocyanin, BioXcell) antibodies were used as control for CD4 depletion. IgG2a (GP120:9674, Genentech; anti-Fluorescein, Absolute Antibody) antibodies were used as control for CD8 and CD20 depletions. IKK RORc Het mice were treated with anti-CD8 or anti-CD20 antibodies between 4.5 months and 6.5 months and with anti-CD4 antibodies between 5.5 months and 6.5 months. IKK RORc KO mice were treated between 3 weeks and 2.3 months with anti-CD8, anti-CD20 antibodies or their combination. IKK RORc KO mice were treated with anti-CD4 antibodies for a long term or short term period (between 3 weeks and 2.3 months, or between 1.6 months and 2.3 months, respectively). To avoid a humoral response against the depleting antibodies, due to prolonged treatment, both anti-CD8 and anti-CD20 antibodies were from mouse host. Anti-CD4 antibodies from rat host were used for a shorter time period.

### Flow cytometry

Flow cytometry data was acquired from fresh or cryopreserved primary splenocytes on a CytoFLEX LX flow cytometer (Beckman Coulter) and analyzed using FCS express software. For detection of cell surface targets, splenocytes were stained for 30 min at 4°C with the following antibodies: APC-CD3, PB-CD8, 488-CD4, and PE-CD19. Non-specific binding of antibodies was blocked using Fc Blocker and rat serum for 20 min at 4°C.

### Liver immune cell isolation, staining and sorting

Immune cells were isolated from 6 months old DEN-injected mice. Immune cells were isolated from 3 IKK RORc Het mice (cells were pooled after isolation), and from 2 IKK RORc KO mice. Frozen sections of one lobe from each mouse were prepared and analyzed immediately by H&E staining, to avoid collecting liver tissue containing lymphoma. Livers were cut to small pieces in 5 ml liver digest medium, squashed with a syringe piston, and incubated at 37⁰c for 30 min. PBS was added and cells were centrifuged at 350g for 5 min at 4⁰c. Further digestion was done with Accumax+DNAse for 10 min at 37⁰c, followed by neutralization with FBS and PBS. Suspension was passed through a 70 µm mesh filter, and two centrifugations at 30g for 5 min at 4⁰c were performed to reduce hepatocytes content. Supernatant was then centrifuged at 350g at 4⁰c to precipitate the immune cells, and the pellet was resuspended in 6ml of 35% Percoll and gently placed on top of 3ml of 75% Percoll. Cells were centrifuged for 20 min at 900g without brake, and the ring of immune cells between the 2 Percoll concentrations was collected into PBS and centrifuged for 8 min at 350g. Red blood cell lysis was performed for 2 min at RT and cells were resuspended in PBS+2%FBS (FACS buffer). Cells were counted using Trypan blue. Three million cells were taken for antibody staining.

Before antibody staining, blocking (FACS buffer, 1%Rat Serum, 1%Fc Blocker) was done for 10 min at 4⁰c. Cells were incubated with primary antibodies (Pacific Blue-CD45 and APC-F4/80) for 20 minutes at 4⁰c, washed with FACS buffer and transferred through a 35μm filter. Viability was assessed by Propidium Iodide which was added to the cells just before sorting.

CD45^+^F4/80^-^ PI^-^ cells were sorted using a BD FACSAria™ III sorter. Sorting was performed using a 85-micron nozzle and threshold was kept in the range of 3000 events/second. F4/80 positive cells (macrophage/Kupffer cells), which constitute a major part of liver immune cells, were excluded from the sort to reduce their amounts from the total CD45 positive cells and enable a better representation of lymphocytes. Cells were sorted into Eppendorf tubes containing 1ml FACS buffer, and then centrifuged without brake. Cells were counted and brought to a concentration of 700 cells/μl.

### Single cell RNA sequencing

Cells were loaded onto a Chromium single-cell 3′ v2 chip (10x Genomics) and libraries were prepared according to the manufacturer’s protocol. Libraries were sequenced by Illumina HiSeq 2500 and the reads were mapped to *mm10* library (ensembl v98). The alignment was performed using *Cellranger* (v6.1.2) (Zheng et al., 2017). Downstream analysis was performed by *Seurat* (v4.0.4) R package (Hao et al., 2021). Cells with a low number of detected cDNA UMI (<1000), a low number of identified genes (<200), or a high percentage of mitochondrial genes (>10%) were filtered out. The standard Seurat single-cell processing pipeline was used for downstream analysis. The parameters for PCA, Cluster determination and UMAP generation were adjusted as necessary to achieve the required resolution. After initial clustering on the cells by UMAP, separate analyses were performed for T cells and for B cells. A total of 3089 cells were analyzed for IKK RORc Het and 7698 cells for IKK RORc KO mice.

### Cytometry by Time-of-Flight (CyTOF)

Immune cells were isolated from 6 months old DEN-injected mice. Immune cells from 4 IKK and from 3 IKK RORc Het mice were isolated and then pooled separately. Immune cells from 3 IKK RORc KO were isolated separately and one lobe was taken for frozen section and H&E staining immediately to avoid collecting liver tissue containing lymphoma. Immune cells were isolated as described for the scRNA-seq method, except for the initial digestion steps: the liver was cut to small pieces and incubated in 5ml Liver digest medium, DNAse + Brefeldin A, and samples were submitted to gentleMACS dissociation. The cells were then incubated at 37⁰c for 15 min and submitted to another step of gentleMACS dissociation.

Isolated cells were washed with Maxpar PBS and stained with Cell-ID Cisplatin viability stain for 2 min. Cells were washed twice with Maxpar Cell Staining Buffer and stained with the extracellular antibodies cocktail at RT for 1 hour (in the presence of Fc blocker and Rat serum). After extracellular staining, cells were washed twice with Maxpar Cell Staining Buffer and fixed overnight at 4⁰c using the Foxp3/Transcription Factor staining buffer set. Centrifugation steps were done at 350g *before* fixation and at 700g *after* fixation to minimize cell loss. Cells were centrifuged, permeabilized and each sample was barcoded for 2 hours at RT using the Cell-ID 20-Plex Pd Barcoding Kit. Cells were washed twice in permeabilization buffer and counted. Equal numbers of cells form each barcoded sample were pooled and then blocked and stained with intracellular antibodies cocktail for 1 hour at RT. Cells were washed with Maxpar Cell Staining Buffer and fixed in 4% PFA. On acquisition day, samples were suspended with 4% PFA supplied with Cell-ID Intercalator-Iridium. Cells were then washed with Maxpar Cell Staining Buffer and then with Maxpar Water and suspended in Maxpar water supplied with 10% EQ Four Element Calibration Beads. Prior to the run on the CyTOF machine, cells were filtered through a 35µM mesh cell strainer. Data was acquired on a CyTOF Helios system (Fluidigm).

CyTOF data underwent the following pre-processing prior to analyses: First, the CyTOF software by Fluidigm was used to normalize and concatenate the acquired data. Then, several gates were applied using the Cytobank platform (Beckman Coulter): First, the event length and the Gaussian residual parameters. Then the beads were gated out using the 140Ce beads channel. Then, live single cells were gated using the cisplatin 195Pt, iridium DNA label in 193Ir, and single cells were gated using the two channels for iridium (Bagwell et al., 2020). Finally, the CyTOF software was used for sample debarcoding. Overall, a total of 133,334 cells were analyzed for each mouse.

### Immunofluorescence

Immunofluorescence was performed on formalin-fixed paraffin-embedded (FFPE) sections. Sections were de-paraffinized in Xylene (twice for 5 min) and rehydrated through a series of 100%, 96% and 80% Ethanol followed by 2 washes in PBS. Antigen retrieval was done in Citrate or Tris buffers in pressure cooker (125⁰c, 3 min). Sections were rinsed twice in PBS and blocked for 2 hours at RT in blocking buffer (2% BSA, 10% Donkey serum, 0.1% Tween). Primary antibody was diluted in blocking buffer and incubated overnight at 4⁰c. Sections were washed in PBS 0.1% Tween, and secondary fluorescent conjugated antibody, diluted in PBS 2% BSA, was incubated at RT for 2 hours. Sections were washed 3 times in PBS and incubated with DAPI for 5 min followed by 2 washing steps. Slides were mounted in Vectashield fluorescent mounting medium. All antibodies and staining conditions can be found in the resource table.

### Immunohistochemistry

Immunohistochemistry was performed on FFPE sections. Sections were de-paraffinized and rehydrated followed by 2 washes in DW. Antigen retrieval was done in Citrate or Tris buffers in pressure cooker (125⁰c, 3 min). Sections were rinsed twice in DW and incubated in 3% Hydrogen Peroxide for 10 min at RT followed by 3 rinses in DW. Sections were washed in wash buffer and blocked with protein block for 2 hours at RT. Sections were incubated with primary antibody diluted in protein block over night at 4⁰c and washed 3 times in wash buffer. Sections were incubated with HRP-conjugated secondary antibodies for 30 min at RT and washed in wash buffer. DAB solution was added for up to 15 min in the dark, and sections were rinsed in tap water. Hematoxylin staining was done for 10 to 60 seconds followed by washes in tap water. Sections were dehydrated and mounted.

### Image analysis

TLSs in IKK mice were defined previously (Finkin et al., 2015) by the presence of B and T cell zones, FDCs and HEVs as well as macrophages and Tregs. In IKK mice, all TLSs contain tumor progenitor cells, and are referred to in the text and figures as “TLSs” unless stated otherwise. In addition to TLSs harboring tumor progenitors, IKK RORc KO mice also displayed TLSs lacking tumor progenitors (indicated in Figures 1 and 2 and their related supplementary Figures). The latter are mostly small in size (less than 150μm in diameter), are also composed of segregated B and T cells zones and were quantified separately from TLSs harboring tumor progenitors. RORc KO mice displayed only TLSs lacking tumor progenitors. TLSs harboring tumor progenitor cells were identified according to H&E staining and validation was done by staining with Sox9 and CD44v6. Tumors were scored as HCC or iCCA according to H&E staining. Validation of tumor type identification was done by IHC for HNF4 and CK19: HCCs were identified by HNF4 positive staining, while iCCAs were defined by lack of HNF4 expression and expression of CK19. In all graphs showing quantification of TLS and tumor numbers per area, all the TLSs and tumors in two sections representing the entire liver were counted, and each data point represents a single mouse. H&E slides were scanned using the Aperio AT2 slide scanner. Aperio ImageScope software was used to quantify the abundance of TLSs and tumors and their diameters. Aperio ImageScope was also used to quantify positive cells in immunohistochemistry. Quantification of staining areas was done using QuPath 0.2.3 software. For Immunofluorescent staining, images were collected on a Nikon Digital Eclipse C1 confocal microscope using a x40 oil lens, converted to tiff format using EZ-C1 software and analyzed using Photoshop software. Cells positive for an antibody staining (or combinations) were scored manually using “Count tool” in a field with identical area. Since the distribution of CD8 and CD4 cells in TLSs is not even, the field was placed in the most crowded area of CD8 or CD4 cells within TLSs. Between 3 and 10 TLSs were scored per mouse. Only TLSs harboring tumor progenitors were analyzed. In all graphs showing quantification of immunostaining, each data point represents an average of several TLSs in a single mouse. The TLSs analyzed by immunostaining for various markers were below 400mm in diameter, since in IKK RORc KO mice there was an increased prevalence of TLSs of this size specifically compared to the control group.

### Statistical analysis

Statistical analysis was performed by an expert statistician. The comparison of a quantitative variable between two independent groups was performed by using the non-parametric Mann-Whitney test. The comparison of a quantitative variable between three or more independent groups was carried out by using the non-parametric Kruskal-Wallis test. The Bonferroni correction of the significance level was applied to Post-Hoc tests. Non-parametric tests were used due to the small sample sizes used in the experiments.

All statistical tests were two-tailed, and a p-value of 5% or less was considered statistically significant. Levels of significance are indicated as *p < 0.05; **p < 0.01; ***p < 0.001. Statistical details for all experiments can be found in the respective figure legends. All graphs show Box plots representing Median, Maximum and Minimum values. Graphs were generated using the Seurat package for R, Cytobank, Excel and GraphPad Prism 6.0 software. Significance is not indicated for scRNA-seq and CyTOF experiments due to small sample sizes, yet similar results obtained using both methods, and validation of some of the results by immunostainings, strengthen their validity.

The graphical abstract was created with BioRender.com.

