## Supplemental Data for "RORc expressing immune cells negatively regulate tertiary lymphoid structure formation and support their pro-tumorigenic functions"

**Supplementary Figure legends**

**Supplementary Figure 1: RORc expressing cells are not necessary for liver TLS neogenesis, but rather negatively regulate TLS formation. (A,B)** Quantification of TLSs harboring tumor progenitors **(A)** and lacking tumor progenitors **(B)** in livers of the indicated experimental groups (the comparison of TLSs harboring tumor progenitors between IKK RORc Het and IKK RORc KO is the same shown in Fig. 1B). H&E stained liver sections demonstrating a TLS harboring **(A)** or lacking **(B)** tumor progenitors are shown to the left of each graph. **(C)** Size distribution of TLSs harboring tumor progenitors in the indicated experimental groups. Each data point represents the number of TLSs of the indicated diameter range per whole liver section of a single mouse. Statistical tests applied: Kruskal-Wallis with Bonferroni correction in A and B; Mann-Whitney in C.

**Supplementary Figure 2: RORc expressing cells are not necessary for liver TLS neogenesis, but rather negatively regulate TLS formation.** Whole slide images of H&E stained livers. Supernumerary TLSs are observed in IKK RORc KO livers in all conditions compared to IKK ROR Het livers.

**Supplementary Figure 3: Cell depletion is the spleen is efficient.** Flow cytometry analyses of splenocytes stained for CD4, CD8, CD3 and CD19 in DEN-injected IKK RORc Het **(A-D)** and IKK RORc KO **(E-J)** mice, following cell depletion regimes as indicated in **A**, **E** and **H**. CD4- **(B)** CD8- **(C)** and CD20- **(D)** depletion in IKK RORc Het mice, as depicted in A. **(F)** CD4 long term and short term depletion in IKK RORc KO mice, as depicted in E. **(G)** CD8, CD20 or combined CD8+CD20 depletion in IKK RORc KO mice, as depicted in E. CD8 depletion **(I)** and CD20 depletion **(J)** in IKK RORc KO mice, as depicted in H. IgG2b or IgG2a isotypes were used as controls for CD4 or CD8 depletions, respectively. Statistical tests applied: Mann-Whitney.

**Supplementary Figure 4: Cell depletion in the liver is efficient at 6 months.** Cell depletion regime in IKK RORc Het **(A-D)** is as indicated in Fig. 2A. Cell depletion regime in IKK RORc KO **(E-G)** is as indicated in Fig. 2H. **(A)** CD4 depletion; TLSs stained for CD4 (red). **(B,E)** CD8 depletion; TLS stained for CD8. **(C,F)** CD20 depletion; TLS stained for CD19. **(D,G)** Quantification of CD19 staining area relative to TLS area following CD20 depletion, corresponding to C,F. Statistical tests applied: Mann-Whitney.

**Supplementary Figure 5: CD4 cells are required for TLS maintenance in IKK RORc Het mice at 6 months.** Depletion regime is as indicated in Fig. 2A. **(A)** Low magnification H&E staining of livers in control and CD4 depleted mice. H&E staining of liver TLS in CD4- **(B)**, CD8- **(C)** and CD20- **(D)** depleted mice, and the corresponding quantification of TLS numbers below and above 400μm in diameter. Statistical tests applied: Mann-Whitney.

**Supplementary Figure 6: Cell depletion in the liver of IKK RORc KO mice is efficient at 2 months.** Cell depletion regime is as indicated in Fig 2E. **(A)** CD4 depletion; TLSs stained for CD4 (red). **(B and C)** CD8, CD20 or combined CD8+CD20 depletion; TLS stained for CD8 **(B)** or for CD19 **(C)**.

**Supplementary Figure 7: CD4 cells are required for TLS maintenance in IKK RORc KO mice at 2 months.** Cell depletion regime is as indicated in Fig. 2E. Low magnification liver H&E staining following control, CD4 long-term or short-term depletion, as indicated. Insets show TLSs.

**Supplementary Figure 8: B cells, but not CD8 cells, are required for TLS formation in IKK RORc KO mice at 6 months.** Depletion regime is as indicated in Fig. 2H. Low and high magnification H&E staining of livers and TLSs in CD8 **(A,B)** or CD20 **(D,E)** depleted mice, and the corresponding quantification of TLS numbers below and above 400μm in diameter **(C,F)**. Statistical tests applied: Mann-Whitney.

**Supplementary Figure 9: B cells are required for TLS neogenesis and tumor progenitor development independently of CD8 cells in IKK RORc KO mice at 2 months.** Cell depletion regime is as indicated in Fig. 2E. Low magnification liver H&E staining following CD8, CD20 or combined CD8+CD20 depletion. Insets show TLSs harboring (top panels) and lacking (lower panels) tumor progenitors.

**Supplementary Figure 10: B cells are required for TLS neogenesis and tumor progenitor development, independently of CD8 cells in IKK RORc KO mice at 2 months.** Cell depletion regime is as indicated in Fig. 2E. Co-immunostaining for the tumor progenitor markers Sox9 (nuclear staining) and CD44v6 (membrane staining) in CD8-, CD20- and combined CD8+CD20-depleted mice. TLSs harboring tumor progenitors stain positively for Sox9 and CD44v6.

**Supplementary Figure 11: Expression of markers in scRNA-seq T cell analysis.** scRNA-seq analysis of $CD45^+$ cells isolated from livers of DEN-injected 6 months old IKK RORc Het mice (pool of 3 mice) and IKK RORc KO mice (2 mice). **(A)** UMAP of $CD45^+$ cells showing cluster identity (left) and the distribution of cells between IKK RORc Het (black) and IKK RORc KO (red) mice (right). **(B)** UMAP restricted to T cells (as in Fig. 4A-C) showing normalized UMI counts of different markers defining T cell subsets and their phenotype (high expression – red, low expression - gray).

**Supplementary Figure 12: Expression of markers in scRNA-seq B cell analysis.** scRNA-seq analysis of $CD45^+$ cells isolated from livers of DEN-injected 6 months old IKK RORc Het mice (pool of 3 mice) and IKK RORc KO mice (2 mice). UMAP restricted to B cells (as in Fig. 4D-G) showing normalized UMI counts of different markers defining B cell subsets and their phenotype (high expression – red, low expression - gray).

**Supplementary Figure 13: Expression of markers in CyTOF analysis of liver immune cells.** Same experiment shown in Fig. 5. **(A)** CyTOF analysis of immune cells isolated from livers of DEN-injected 6 months old IKK RORc Het and IKK RORc KO mice. Expression of different markers defining cell subsets and their phenotype is shown on tSNE plots (high expression – red, low expression - blue). **(B)** Immunofluorescent staining of 6 months old DEN-injected IKK RORc Het (top) and IKK RORc KO (bottom) TLSs for CD8 (red) and CD103 (green). Double positive cells are marked by arrowheads. Insets correspond to areas in white boxes. **(C)** Quantification of the staining in B. Statistical test applied: Mann-Whitney.

**Supplementary Figure 14: RORcECs restrict CD8 effector and memory cells with a progenitor phenotype.** Same experiment shown in Fig. 5. **(A)** Expression of indicated genes on tSNE plots (high expression - red, low expression - blue). **(B-G)** Comparisons of specific cell percentages. Note that the denominator population is different in different graphs, as indicated. **(H)** Histograms of cell counts as a function of signal intensity in IKK RORc Het (2 lower histograms) vs. IKK RORc KO (3 upper histograms) for Eomes and Tbet in the $PD1^-$ expressing $CD8_{EFF}$ and $CD8_{MEM}$ populations. GzmB and pS6 expression intensity are shown in the $PD1^{high}$ $CD8_{EFF}$

population and Gata3 expression intensity is shown in the $CD8_{MEM}$ population (high expression - yellow, low expression - black).

**Supplementary Figure 15: Total CD4 cells, Tregs and MDSCs numbers in TLSs are not significantly affected in IKK RORc KO mice. (A)** Triple immunofluorescent staining for CD4, PD1 and Ki67 in TLSs. Arrows point to $CD4^+$ $PD1^+$ cells, and arrowheads to $CD4^+$ $PD1^+$ $Ki67^+$ cells. Insets correspond to areas in white boxes. In merged panels, cells co-expressing CD4 (red) and PD1 (blue) are seen in pink. **(B)** Coimmunofluorescent staining for PD1 and Foxp3. Insets correspond to areas in white boxes. **(C)** Number of cells expressing CD4, PD1 or Ki67 (left) and their combinations (middle), and Foxp3 and PD1 and their combination (right) within TLSs. **(D)** Staining for the MDSC marker Ly6G. Insets correspond to areas in black boxes. Statistical test applied: Mann-Whitney.

**Supplementary Figure 16: CD4 depletion in 6 months old DEN-injected IKK RORc Het mice.** Depletion regime is as indicated in Fig. 8A. Low and high magnification images of H&E stained liver sections. Two sections per mouse, representing the entire liver are shown **(A, A'** and **B, B')**. HCCs are circled in light blue and iCCAs in red. Tumors indicated by arrowheads in A and B',B are shown in higher magnification in **C**,**F** and **D**,**G**, respectively. iCCA **(E)** and HCC **(H)** numbers below and above 1000μm in diameter. Statistical test applied: Mann-Whitney.

**Supplementary Figure 17: CD8 depletion in 6 months old DEN-injected IKK RORc Het mice.** Depletion regime is as indicated in Fig. 8A. Low and high magnification images of H&E stained liver sections. Two sections per mouse, representing the entire liver are shown **(A, A' and B, B')**. HCCs are circled in light blue and iCCAs in red. Tumors indicated by arrowheads in A, A' and B',B are shown in higher magnification in **C**,**F** and **D**,**G**, respectively. iCCA **(E)** and HCC **(H)** numbers below and above 1000μm in diameter. Statistical test applied: Mann-Whitney.

**Supplementary Figure 18: B cells limit iCCA formation in 6 months old DEN-injected IKK RORc Het mice.** Depletion regime is as indicated in Fig. 8A. Low and high magnification images of H&E stained liver sections. Two sections per mouse, representing the entire liver are shown **(A, A' and B, B')**. HCCs are circled in light blue and iCCAs in red. Tumors indicated by arrowheads in A and

B’,B are shown in higher magnification in **C**,**F** and **D**,**G**, respectively. iCCA **(E)** and HCC **(H)** numbers below and above 1000μm in diameter. Statistical test applied: Mann-Whitney.

**Supplementary Figure 19: CD20 depletion reduces the abundance of plasma cells, IgG and IgA in TLSs of IKK RORc Het mice.** Depletion regime is as indicated in Fig. 8A. Staining of serial sections for CD138 **(A)**, IgG **(B)** and IgA **(C)** in TLSs of control and CD20 depleted 6 months old DEN-injected IKK RORc Het mice.

**Supplementary Figure 20: CD8 and CD20 depletion in 6 months old DEN-injected IKK RORc KO mice.** Cell depletion timeline in DEN-injected IKK RORc KO mice **(A)**. HCC number in CD8- **(B)** and CD20- depleted **(C)** IKK RORc KO mice. iCCA number in CD8- **(D)** and CD20- **(E)** depleted IKK RORc KO mice. Ratio of iCCA numbers to TLS numbers in CD8- **(F)** and CD20- **(G)** depleted IKK RORc KO mice. **(H)** Qualitative quantification of the percent of TLSs containing high and medium density of CD138$^{+}$ plasma cells in CD20 depleted IKK RORc KO mice. **(I)** Quantification of IgA staining area relative to TLS area in CD20 depleted IKK RORc KO mice. Each dot represents an average of all TLSs in a mouse (I). Statistical test applied: Mann-Whitney.

**Supplementary Figure 21: CD8 depletion in 6 months old DEN-injected IKK RORc KO mice.** Depletion regime is as indicated in Fig. S20A. Low and high magnification images of H&E stained liver sections. Two sections per mouse, representing the entire liver are shown **(A, A’** and **B, B’)**. HCCs are circled in light blue and iCCAs in red. Tumors indicated by arrowheads in A, A’ and B,B’ are shown in higher magnification in **C**,**F** and **D**,**G**, respectively. iCCA **(E)** and HCC **(H)** numbers below and above 1000μm in diameter. Statistical test applied: Mann-Whitney.

**Supplementary Figure 22: CD20 depletion in 6 months old DEN-injected IKK RORc KO mice.** Depletion regime is as indicated in Fig. S20A. Low and high magnification images of H&E stained liver sections. Two sections per mouse, representing the entire liver are shown **(A, A’ and B, B’)**. HCCs are circled in light blue and iCCAs in red. Tumors indicated by arrowheads in A and B are shown in higher magnification in **C**,**F** and **D**,**G**, respectively. iCCA **(E)** and HCC **(H)** numbers below and above 1000μm in diameter. Statistical test applied: Mann-Whitney.

**Supplementary Figure 23: CD20 depletion increases the abundance of plasma cells, IgG and IgA in TLSs of 6 months old DEN-injected IKK RORc KO mice.** Depletion regime is as indicated in Fig. S20A. Staining of serial sections for CD138 **(A)**, IgG **(B)** and IgA **(C)** in TLSs of controls and CD20 depleted 6 months old IKK RORc KO mice.

**Figure S1.**

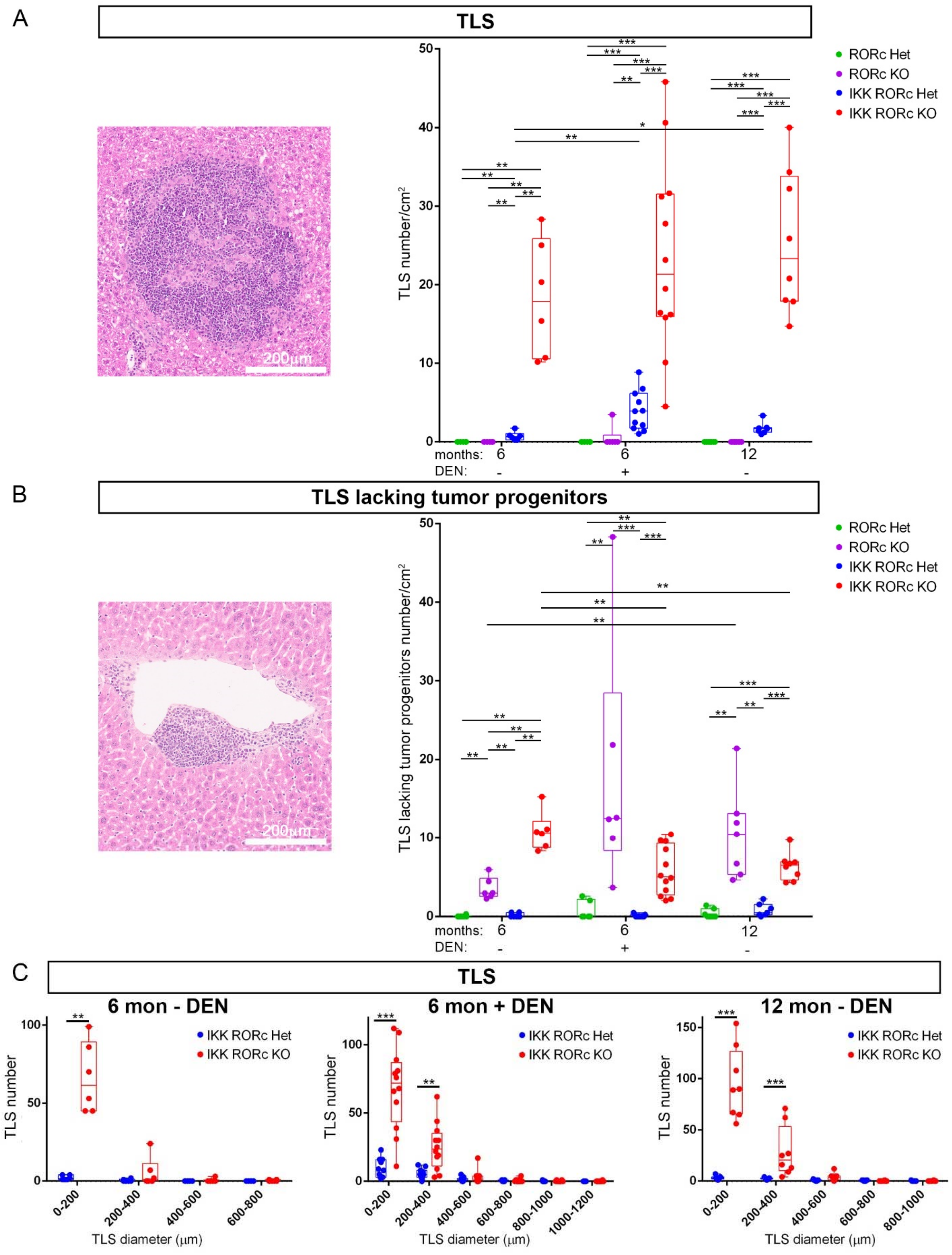
A
TLS
B
TLS lacking tumor progenitors
C
TLS
6 mon - DEN
6 mon + DEN
12 mon - DEN
RORc Het
RORc KO
IKK RORc Het
IKK RORc KO
TLS number/cm2
TLS lacking tumor progenitors number/cm2
months: 6 6 12
DEN: - + -
200μm
TLS number
TLS diameter (μm)

**Figure S2.**

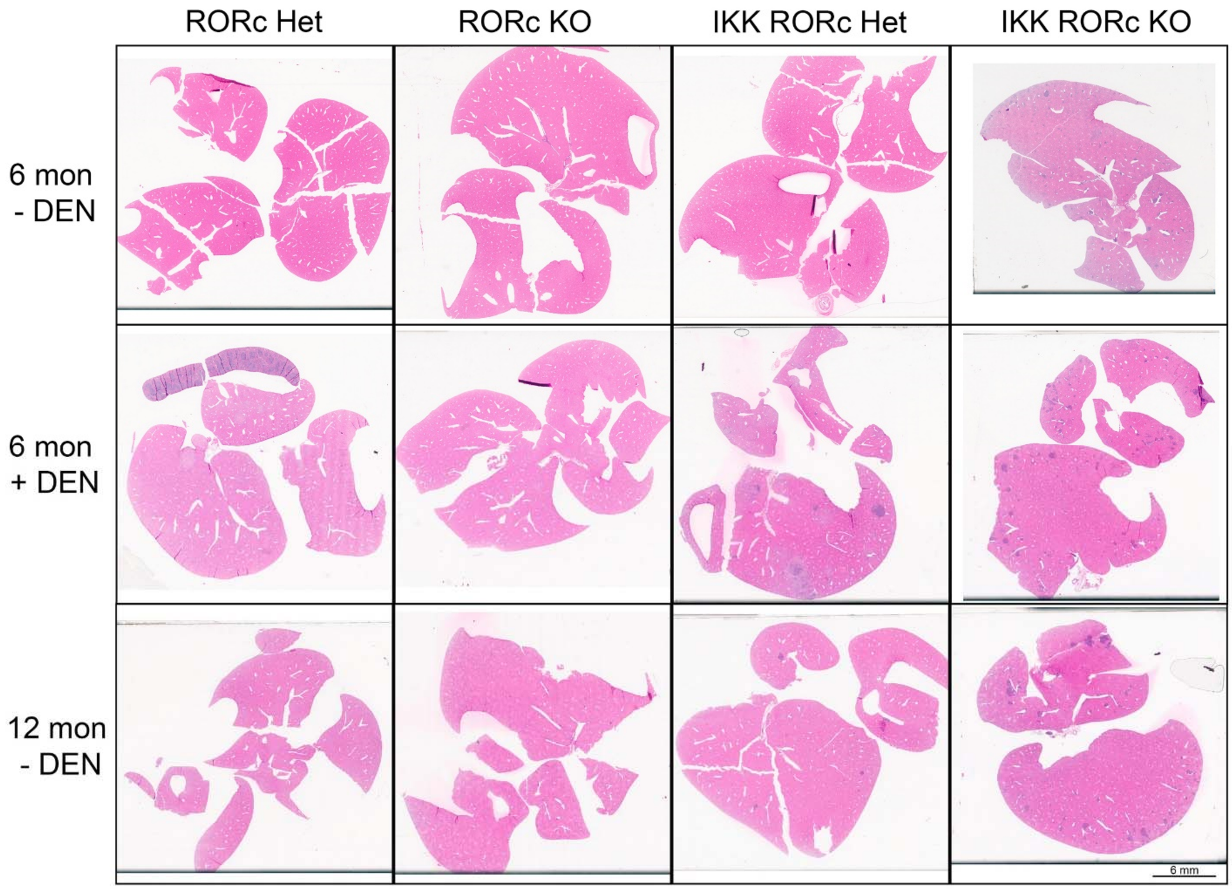

RORc Het
RORc KO
IKK RORc Het
IKK RORc KO
6 mon
- DEN
6 mon
+ DEN
12 mon
- DEN
6 mm

**Figure S3.**

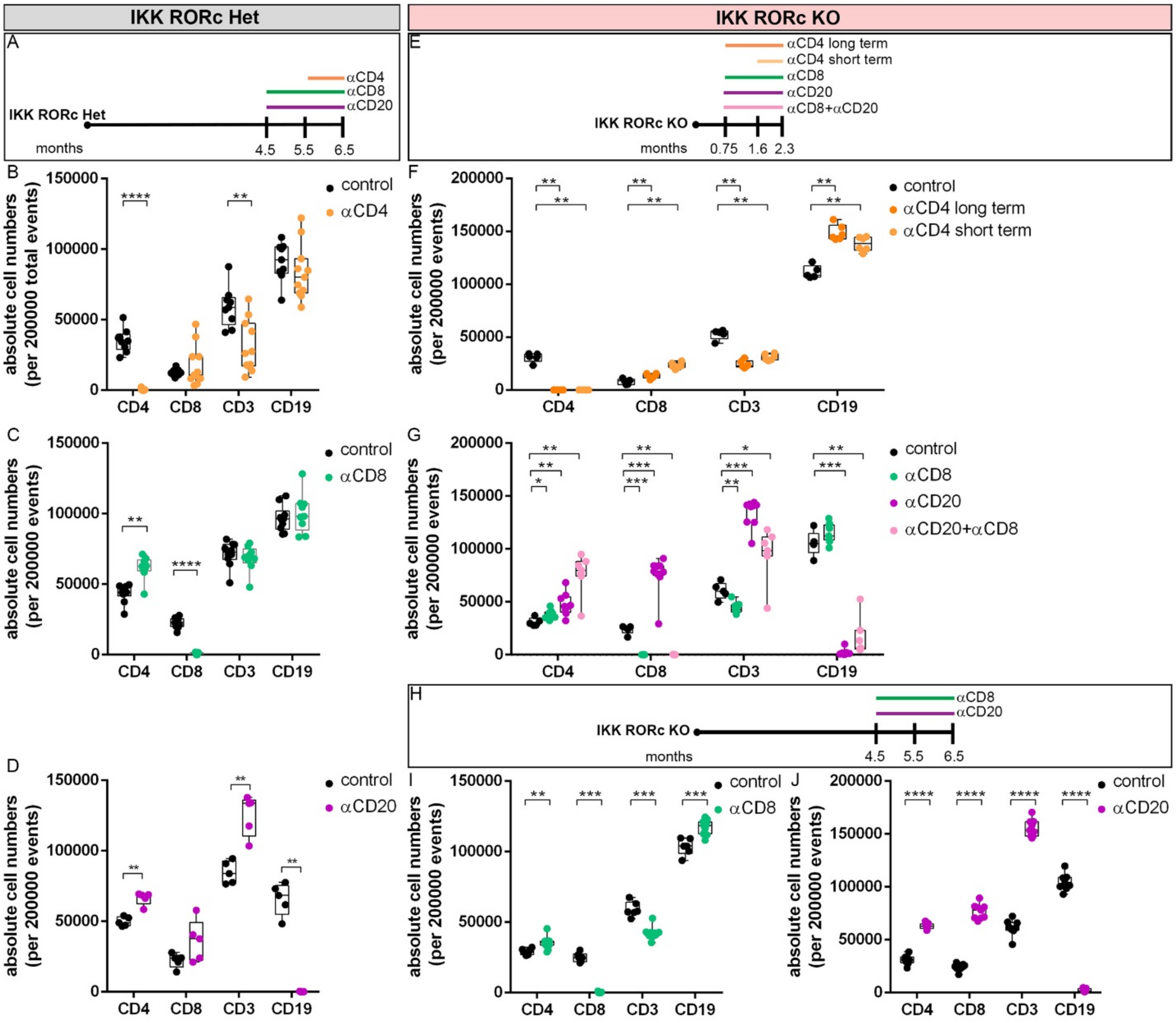
IKK RORc Het
IKK RORc KO
A
αCD4
αCD8
αCD20
IKK RORc Het
months
4.5
5.5
6.5
E
αCD4 long term
αCD4 short term
αCD8
αCD20
αCD8+αCD20
IKK RORc KO
months
0.75
1.6
2.3
B
absolute cell numbers (per 200000 total events)
control
αCD4
CD4
CD8
CD3
CD19
F
absolute cell numbers (per 200000 events)
control
αCD4 long term
αCD4 short term
C
control
αCD8
G
control
αCD8
αCD20
αCD20+αCD8
H
αCD8
αCD20
IKK RORc KO
months
4.5
5.5
6.5
D
control
αCD20
I
control
αCD8
J
control
αCD20

**Figure S4.**

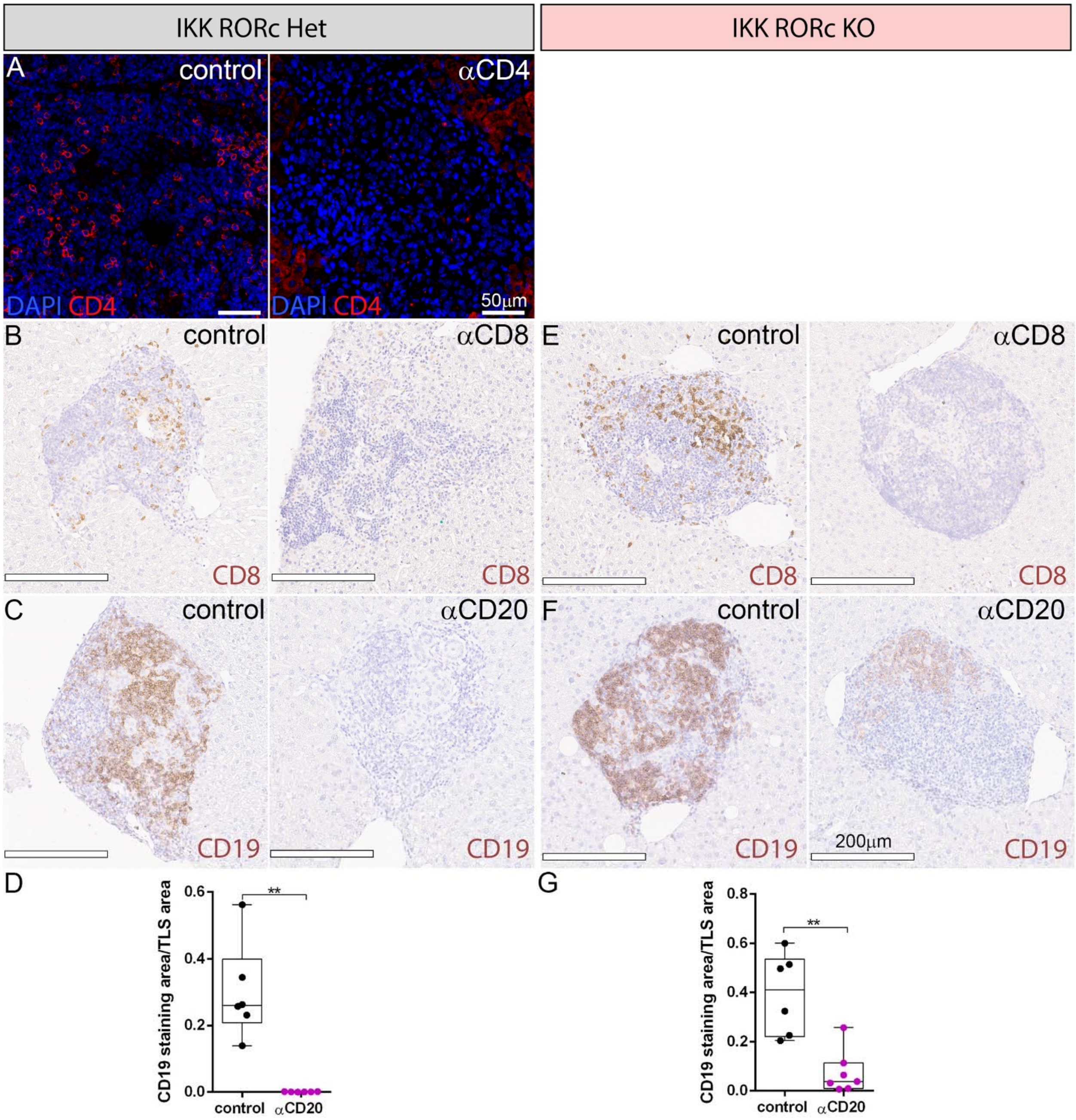

IKK RORc Het
IKK RORc KO
A
control
αCD4
DAPI CD4
DAPI CD4
50μm
B
control
αCD8
CD8
CD8
E
control
αCD8
CD8
CD8
C
control
αCD20
CD19
CD19
F
control
αCD20
CD19
200μm
CD19
D
CD19 staining area/TLS area
0.6
0.4
0.2
0.0
**
control
αCD20
G
CD19 staining area/TLS area
0.8
0.6
0.4
0.2
0.0
**
control
αCD20

**Figure S5.**

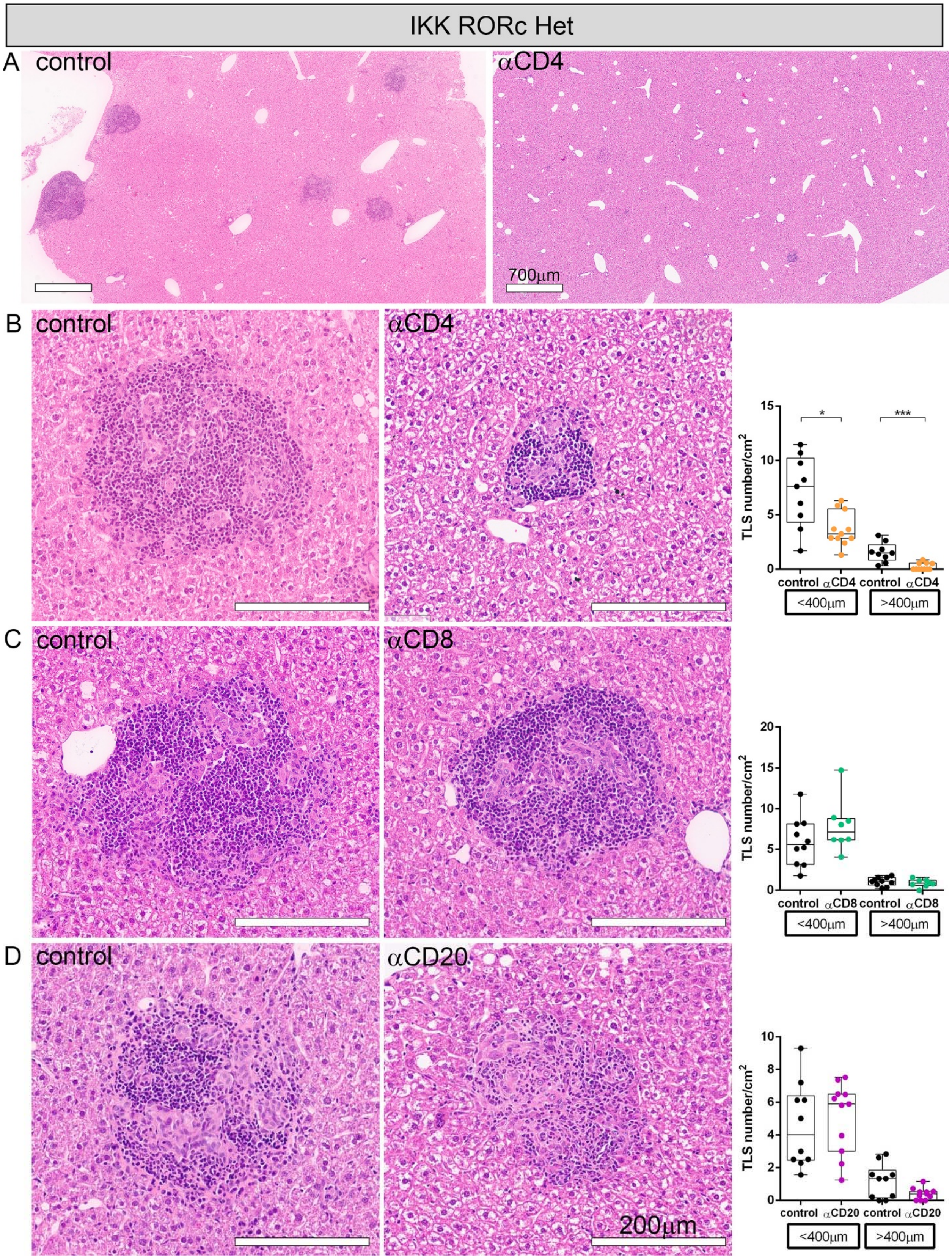

IKK RORc Het
A control
αCD4
700μm
B control
αCD4
TLS number/cm²
*
***
control αCD4 control αCD4
<400μm >400μm
C control
αCD8
TLS number/cm²
control αCD8 control αCD8
<400μm >400μm
D control
αCD20
200μm
TLS number/cm²
control αCD20 control αCD20
<400μm >400μm

**Figure S6.**

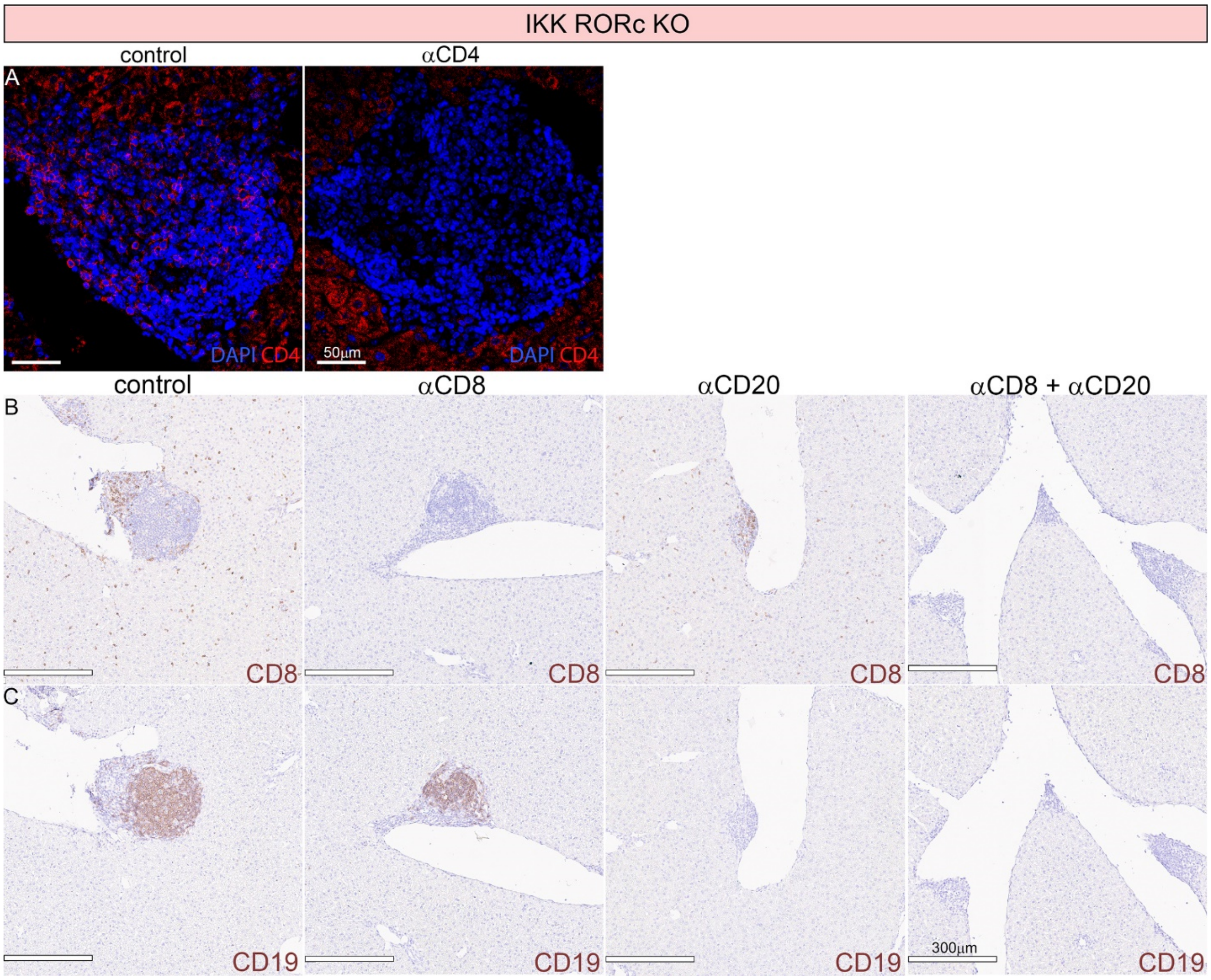

IKK RORc KO
control
αCD4
A
DAPI CD4
50μm
DAPI CD4
control
αCD8
αCD20
αCD8 + αCD20
B
CD8
CD8
CD8
CD8
C
CD19
CD19
CD19
300μm
CD19

**Figure S7.**

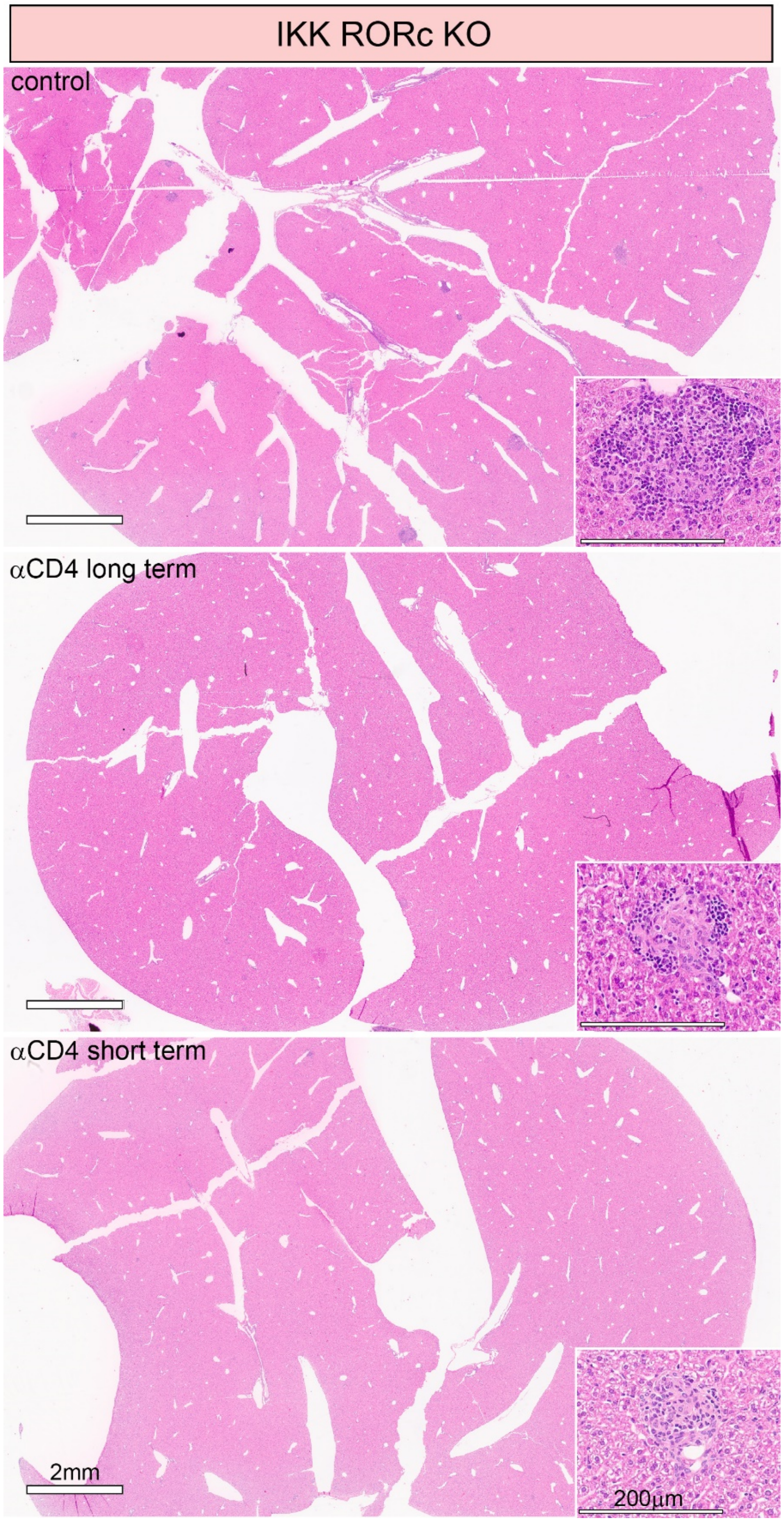

**Figure S8.**

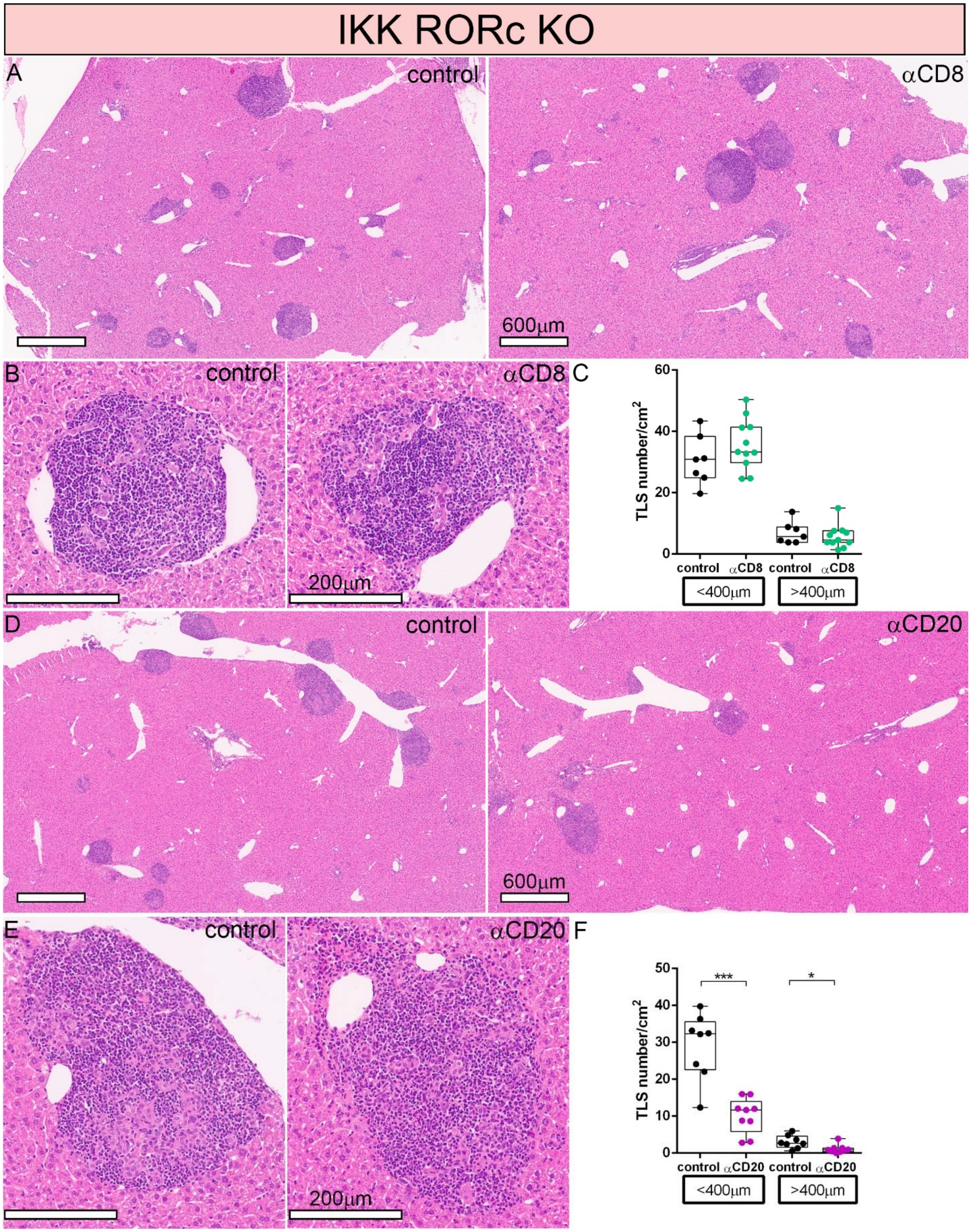
IKK RORc KO
A
control
αCD8
600μm
B
control
αCD8
200μm
C
TLS number/cm²
60
40
20
0
control
αCD8
control
αCD8
<400μm
>400μm
D
control
αCD20
600μm
E
control
αCD20
200μm
F
TLS number/cm²
50
40
30
20
10
0
***
*
control
αCD20
control
αCD20
<400μm
>400μm

**Figure S9.**

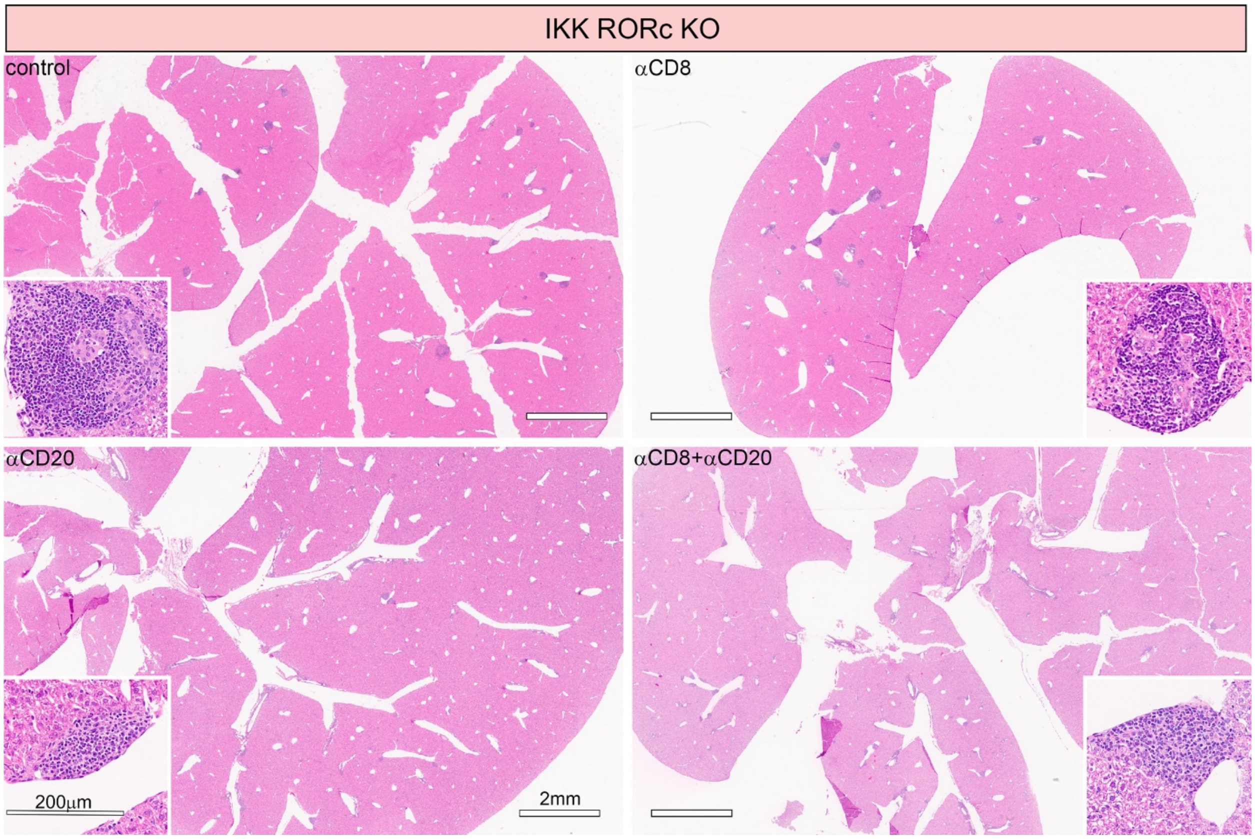

**Figure S10.**

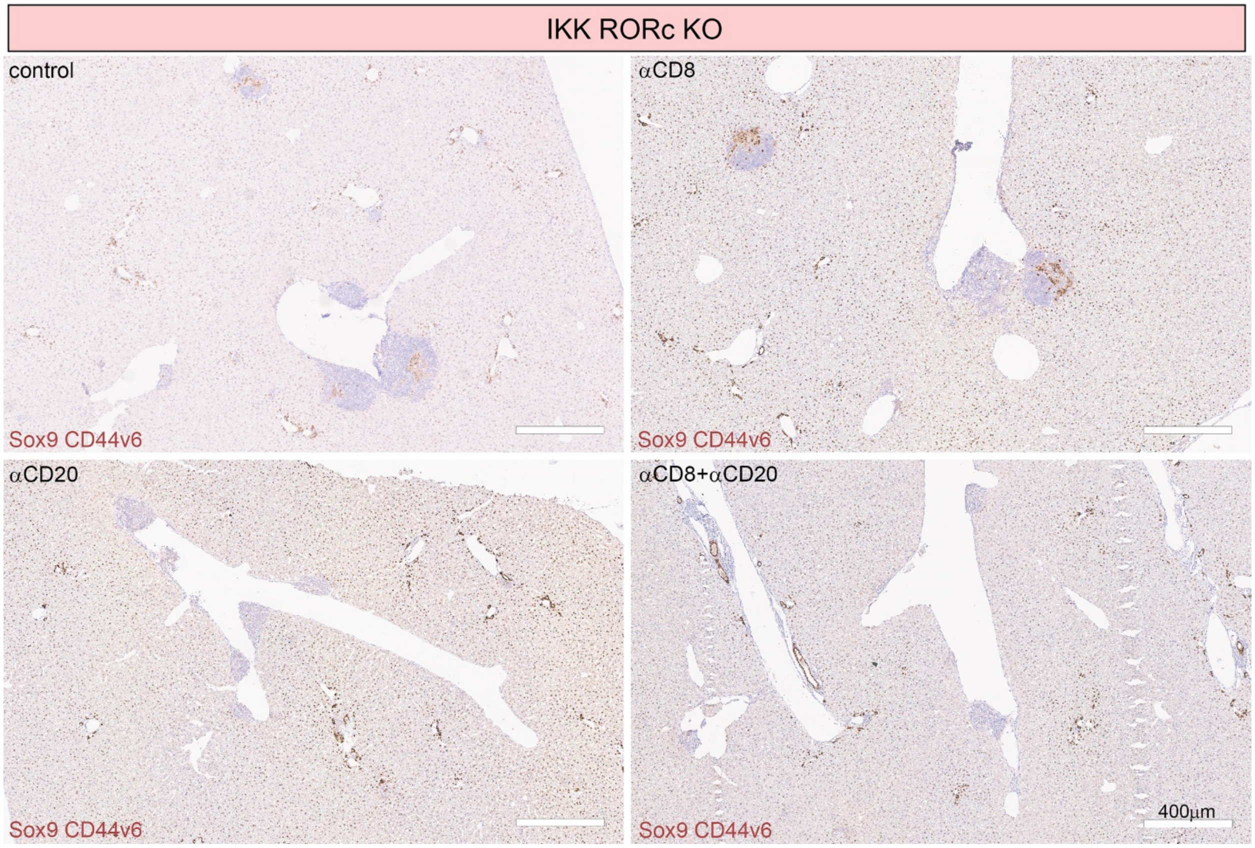
IKK RORc KO
control
Sox9 CD44v6
αCD8
Sox9 CD44v6
αCD20
Sox9 CD44v6
αCD8+αCD20
400μm
Sox9 CD44v6

**Figure S11.**

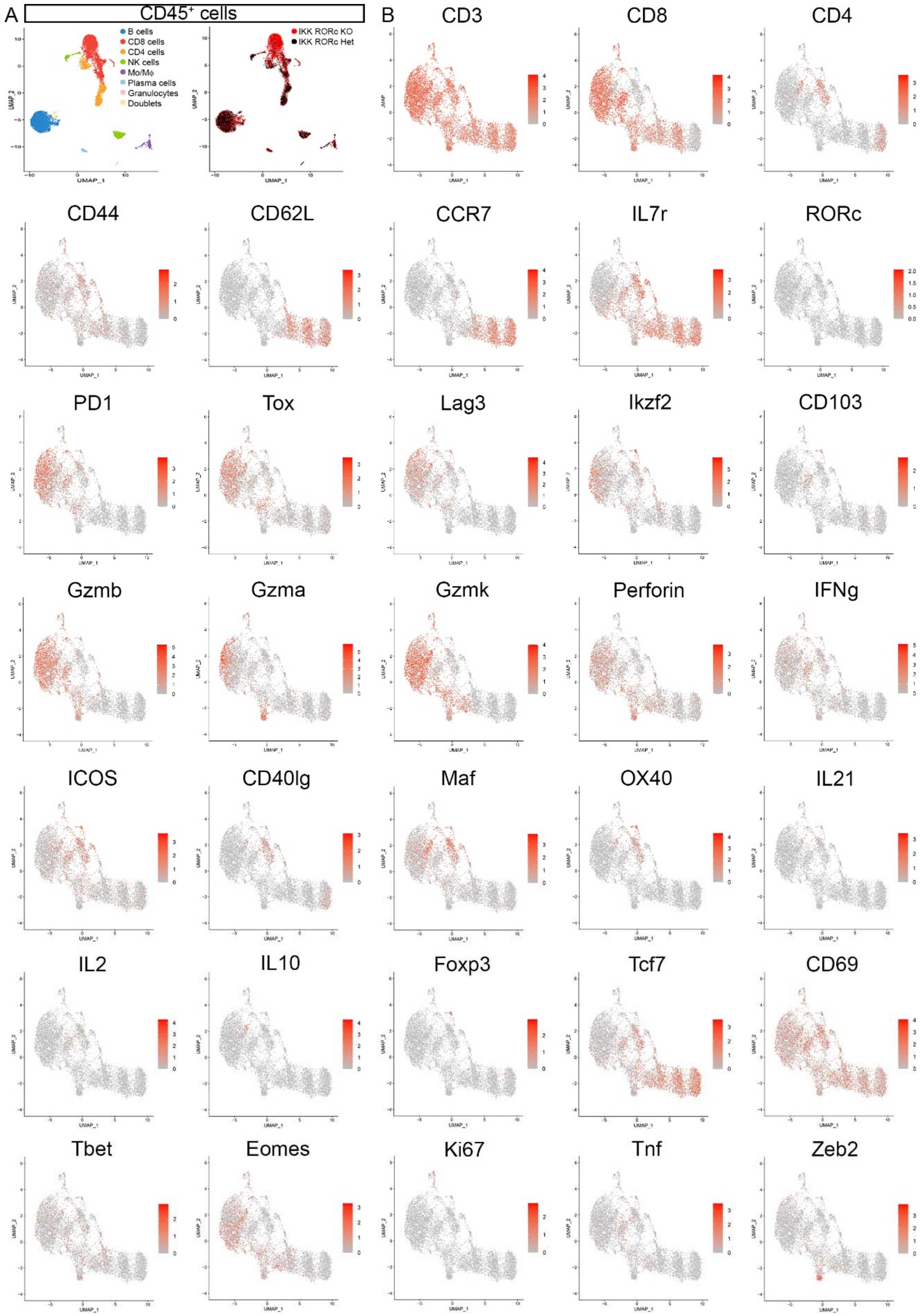
A
CD45+ cells
B cells
CD8 cells
CD4 cells
NK cells
Mo/Mϕ
Plasma cells
Granulocytes
Doublets
IKK RORc KO
IKK RORc Het
B
CD3
CD8
CD4
CD44
CD62L
CCR7
IL7r
RORc
PD1
Tox
Lag3
Ikzf2
CD103
Gzmb
Gzma
Gzmk
Perforin
IFNg
ICOS
CD40lg
Maf
OX40
IL21
IL2
IL10
Foxp3
Tcf7
CD69
Tbet
Eomes
Ki67
Tnf
Zeb2

**Figure S12.**

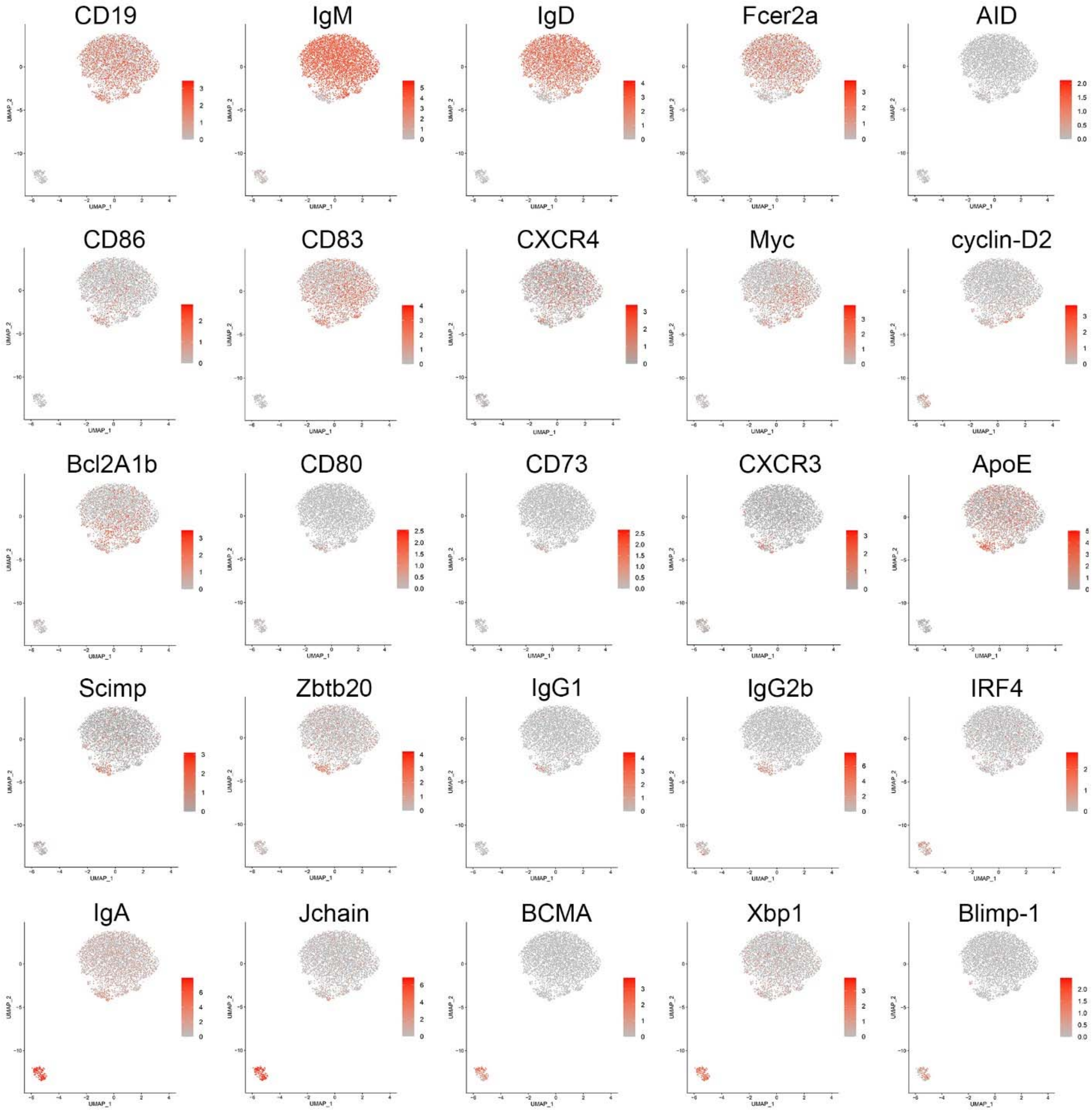
CD19
IgM
IgD
Fcer2a
AID
CD86
CD83
CXCR4
Myc
cyclin-D2
Bcl2A1b
CD80
CD73
CXCR3
ApoE
Scimp
Zbtb20
IgG1
IgG2b
IRF4
IgA
Jchain
BCMA
Xbp1
Blimp-1
UMAP_1
UMAP_2

**Figure S13.**

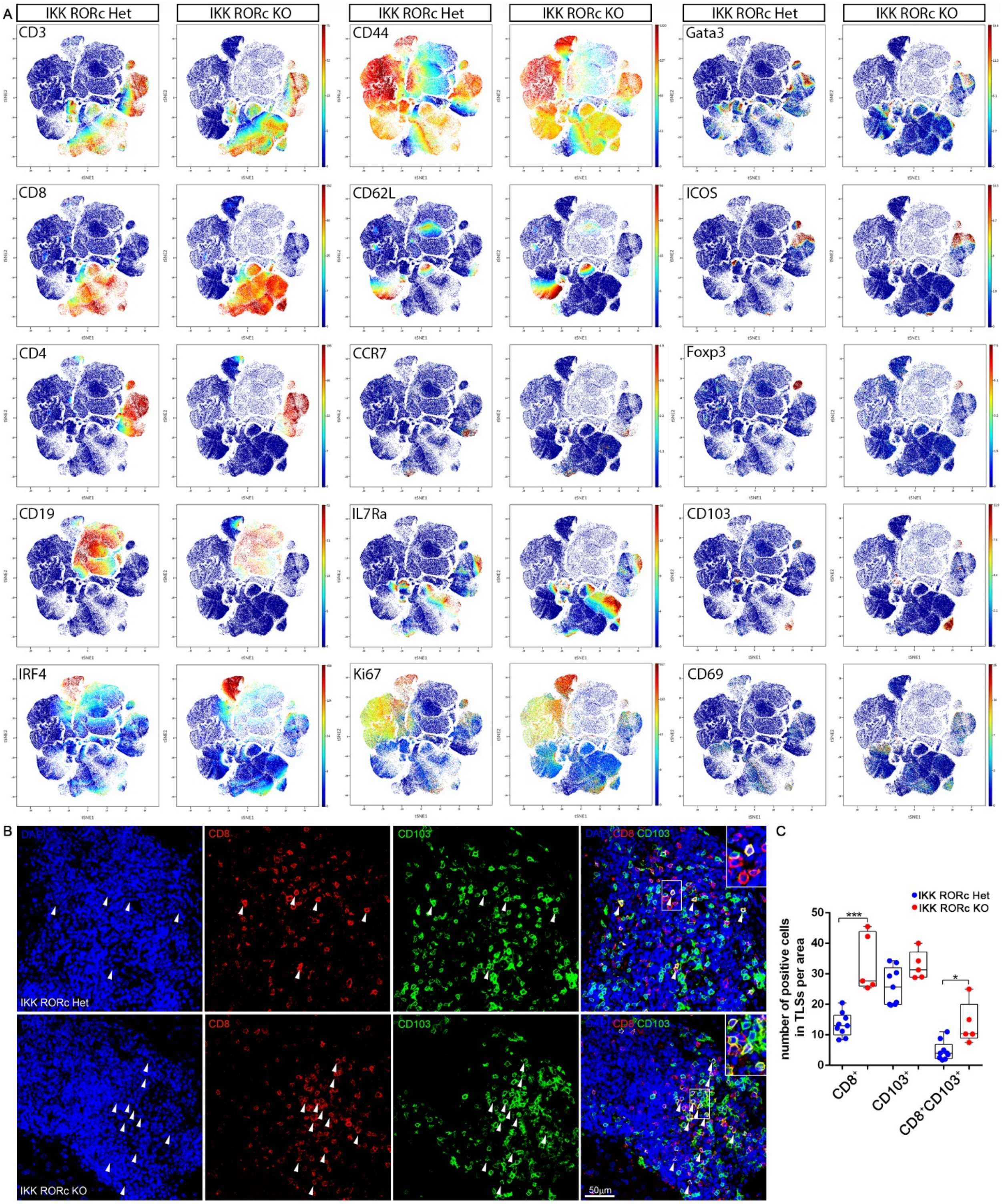
A
IKK RORc Het
IKK RORc KO
IKK RORc Het
IKK RORc KO
IKK RORc Het
IKK RORc KO
CD3
CD44
Gata3
CD8
CD62L
ICOS
CD4
CCR7
Foxp3
CD19
IL7Ra
CD103
IRF4
Ki67
CD69
B
DAPI
CD8
CD103
DAPI CD8 CD103
IKK RORc Het
IKK RORc KO
50μm
C
IKK RORc Het
IKK RORc KO
number of positive cells in TLSs per area
50
40
30
20
10
0
***
*
CD8+
CD103+
CD8+CD103+

**Figure S14.**

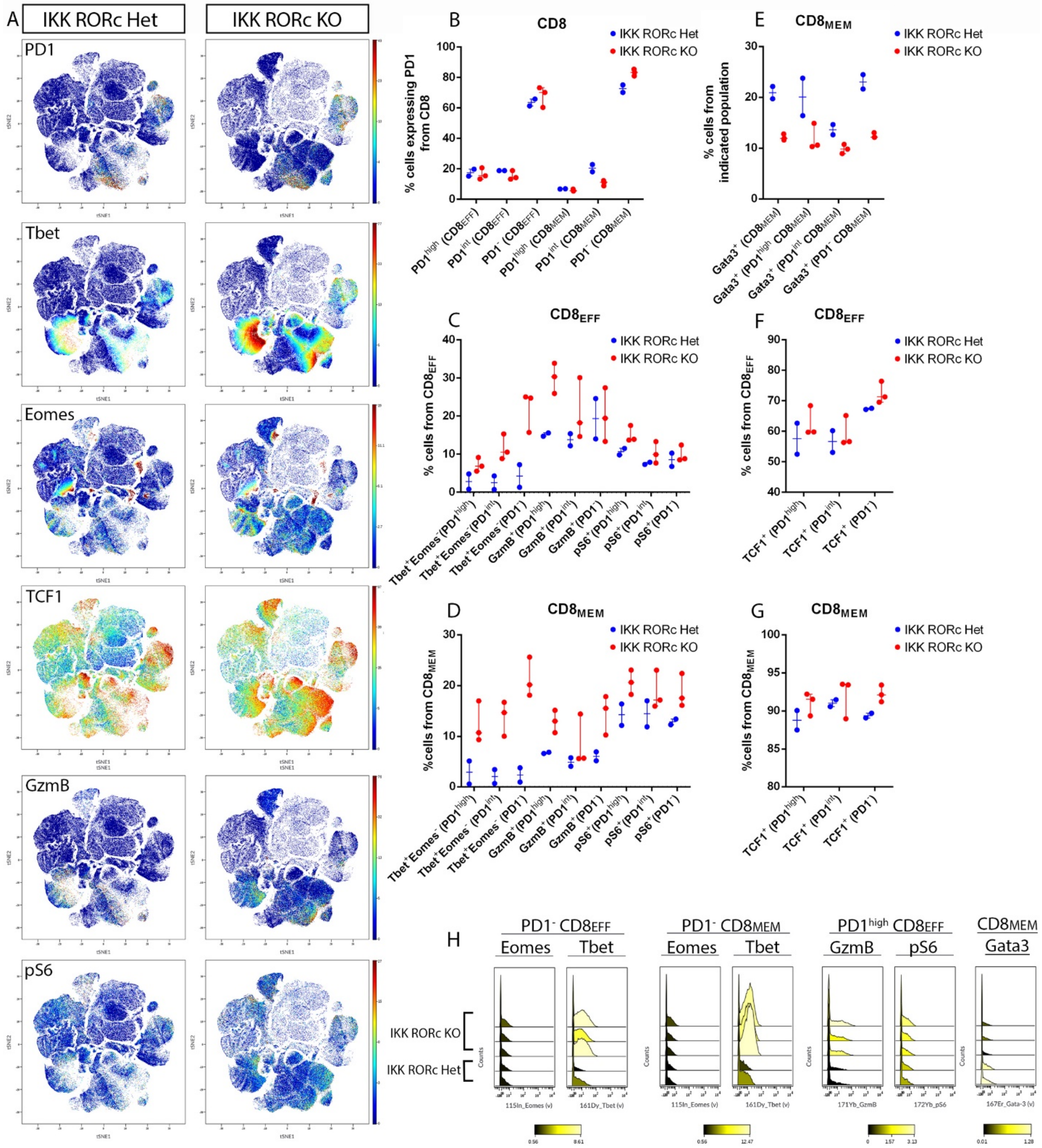
A
IKK RORc Het
IKK RORc KO
PD1
Tbet
Eomes
TCF1
GzmB
pS6
B
CD8
IKK RORc Het
IKK RORc KO
% cells expressing PD1 from CD8
PD1high (CD8EFF)
PD1int (CD8EFF)
PD1- (CD8EFF)
PD1high (CD8MEM)
PD1int (CD8MEM)
PD1- (CD8MEM)
E
CD8MEM
% cells from indicated population
Gata3+ (CD8MEM)
Gata3+ (PD1high CD8MEM)
Gata3+ (PD1int CD8MEM)
Gata3+ (PD1- CD8MEM)
C
CD8EFF
% cells from CD8EFF
Tbet+Eomes-(PD1high)
Tbet+Eomes-(PD1int)
Tbet+Eomes (PD1-)
GzmB+(PD1high)
GzmB+(PD1int)
GzmB+(PD1-)
pS6+(PD1high)
pS6+(PD1int)
pS6+(PD1-)
F
CD8EFF
TCF1+ (PD1high)
TCF1+ (PD1int)
TCF1+ (PD1-)
D
CD8MEM
%cells from CD8MEM
G
CD8MEM
% cells from CD8MEM
H
PD1- CD8EFF
PD1- CD8MEM
PD1high CD8EFF
CD8MEM
Eomes
Tbet
GzmB
pS6
Gata3
IKK RORc KO
IKK RORc Het
Counts
115In_Eomes (v)
161Dy_Tbet (v)
171Yb_GzmB
172Yb_pS6
167Er_Gata-3 (v)

**Figure S15.**

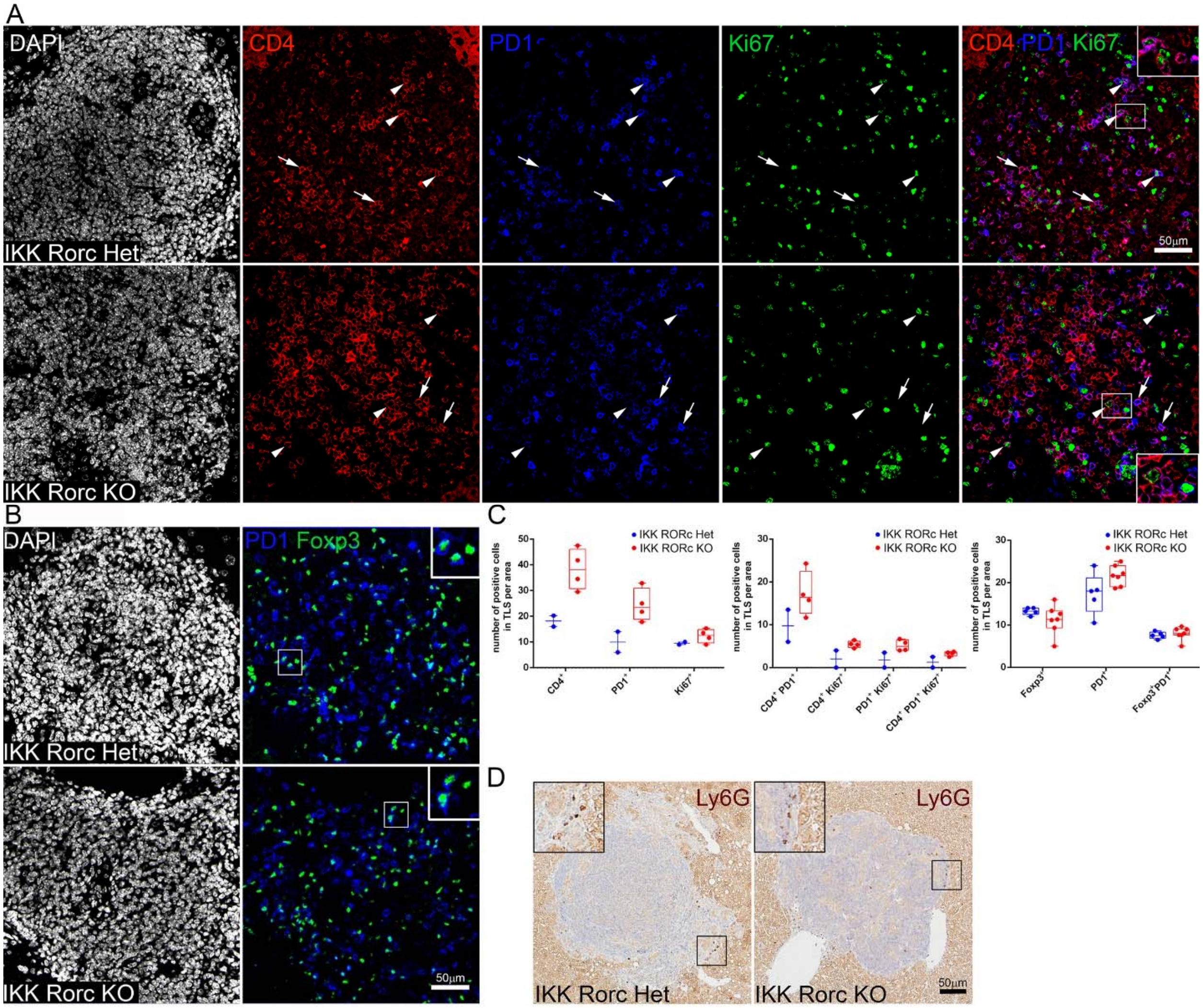
A
DAPI
CD4
PD1
Ki67
CD4 PD1 Ki67
50μm
IKK Rorc Het
IKK Rorc KO
B
DAPI
PD1 Foxp3
IKK Rorc Het
IKK Rorc KO
50μm
C
IKK RORc Het
IKK RORc KO
number of positive cells in TLS per area
CD4+
PD1+
Ki67+
CD4+ PD1+
CD4+ Ki67+
PD1+ Ki67+
CD4+ PD1+ Ki67+
Foxp3+
Foxp3+PD1+
D
Ly6G
IKK Rorc Het
IKK Rorc KO
50μm

**Figure S16**.

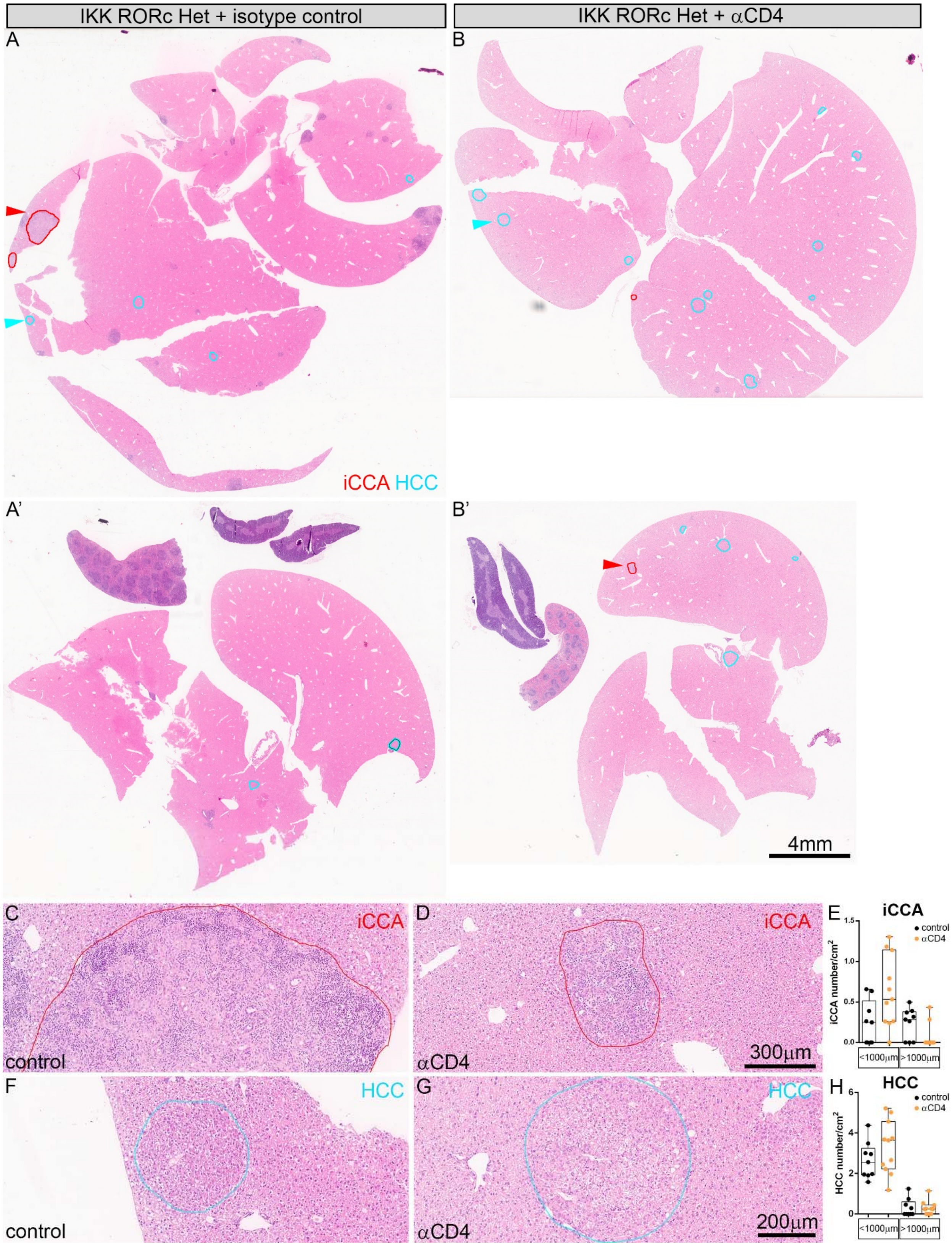
IKK RORc Het + isotype control
IKK RORc Het + αCD4
A
B
iCCA HCC
A'
B'
4mm
C
iCCA
control
D
iCCA
αCD4
300μm
E
iCCA
control
αCD4
iCCA number/cm²
<1000μm
>1000μm
F
HCC
control
G
HCC
αCD4
200μm
H
HCC
control
αCD4
HCC number/cm²
<1000μm
>1000μm

**Figure S17.**

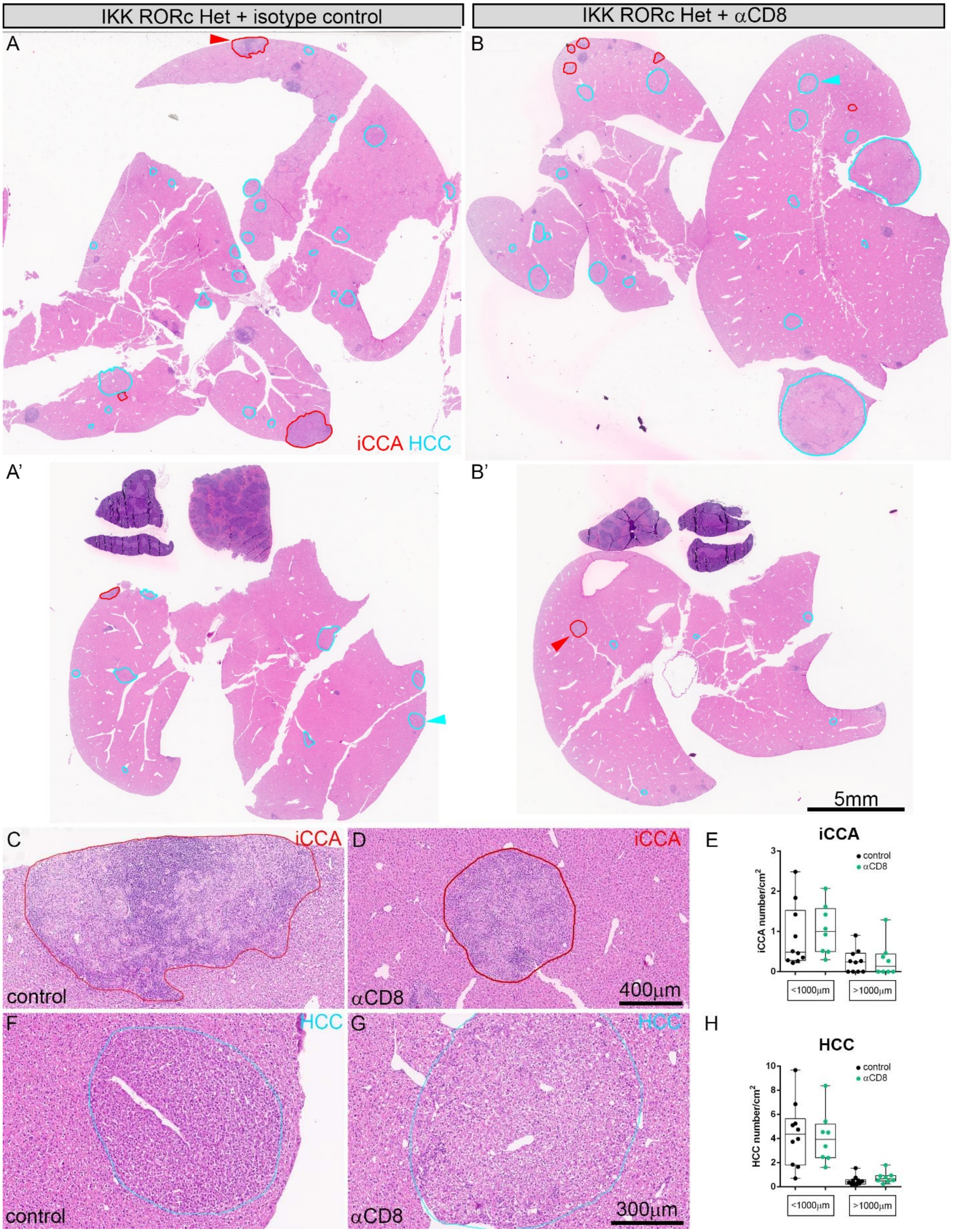
IKK RORc Het + isotype control
IKK RORc Het + αCD8
A
B
iCCA HCC
A'
B'
5mm
C
iCCA
control
D
iCCA
αCD8
400μm
E
iCCA
control
αCD8
iCCA number/cm2
<1000μm
>1000μm
F
HCC
control
G
HCC
αCD8
300μm
H
HCC
control
αCD8
HCC number/cm2
<1000μm
>1000μm

**Figure S18.**

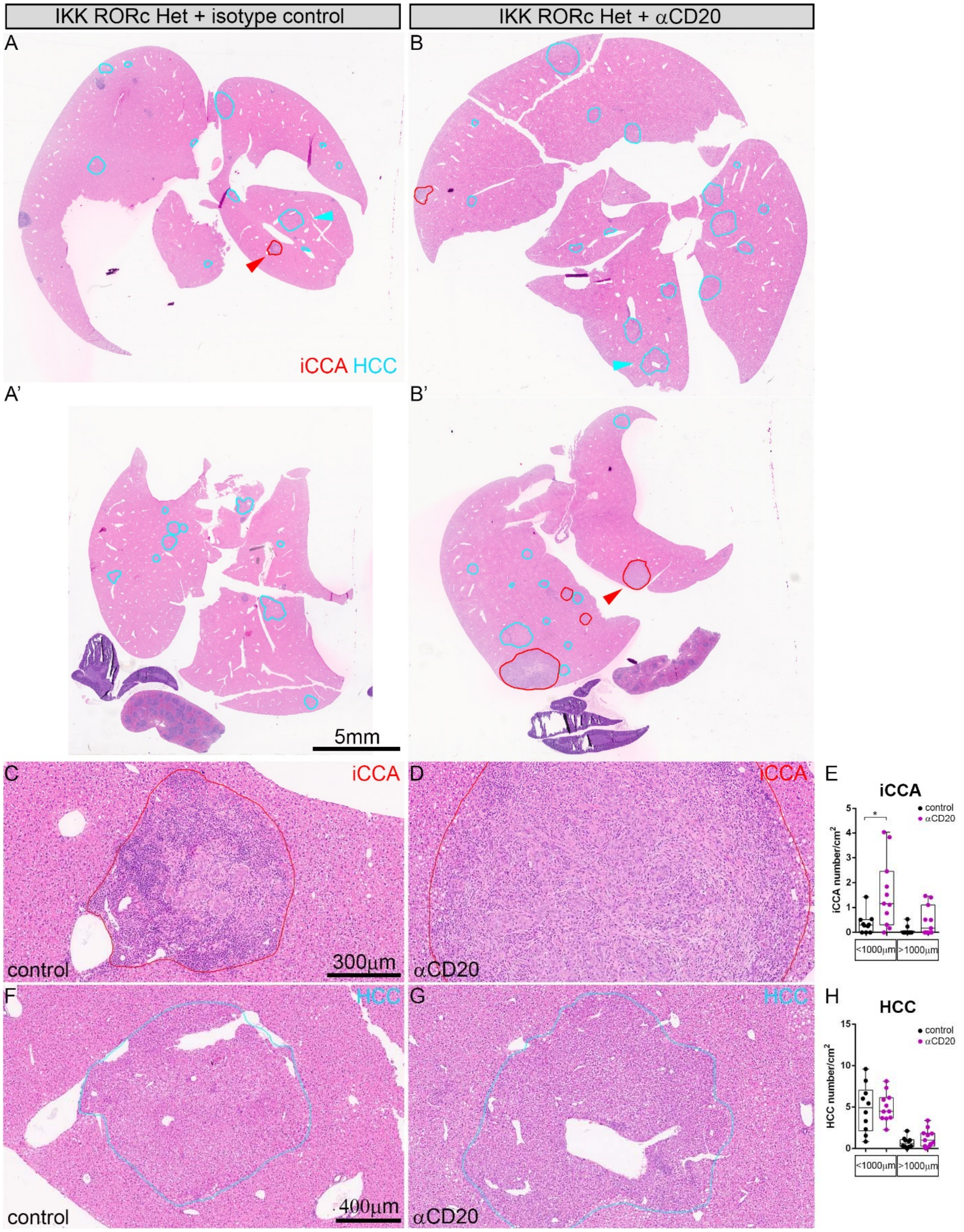
IKK RORc Het + isotype control
IKK RORc Het + αCD20
A
B
iCCA HCC
A’
B’
5mm
C
iCCA
control
300µm
D
iCCA
αCD20
E
iCCA
control
αCD20
iCCA number/cm²
<1000µm
>1000µm
F
HCC
control
400µm
G
HCC
αCD20
H
HCC
control
αCD20
HCC number/cm²
<1000µm
>1000µm

**Figure S19.**

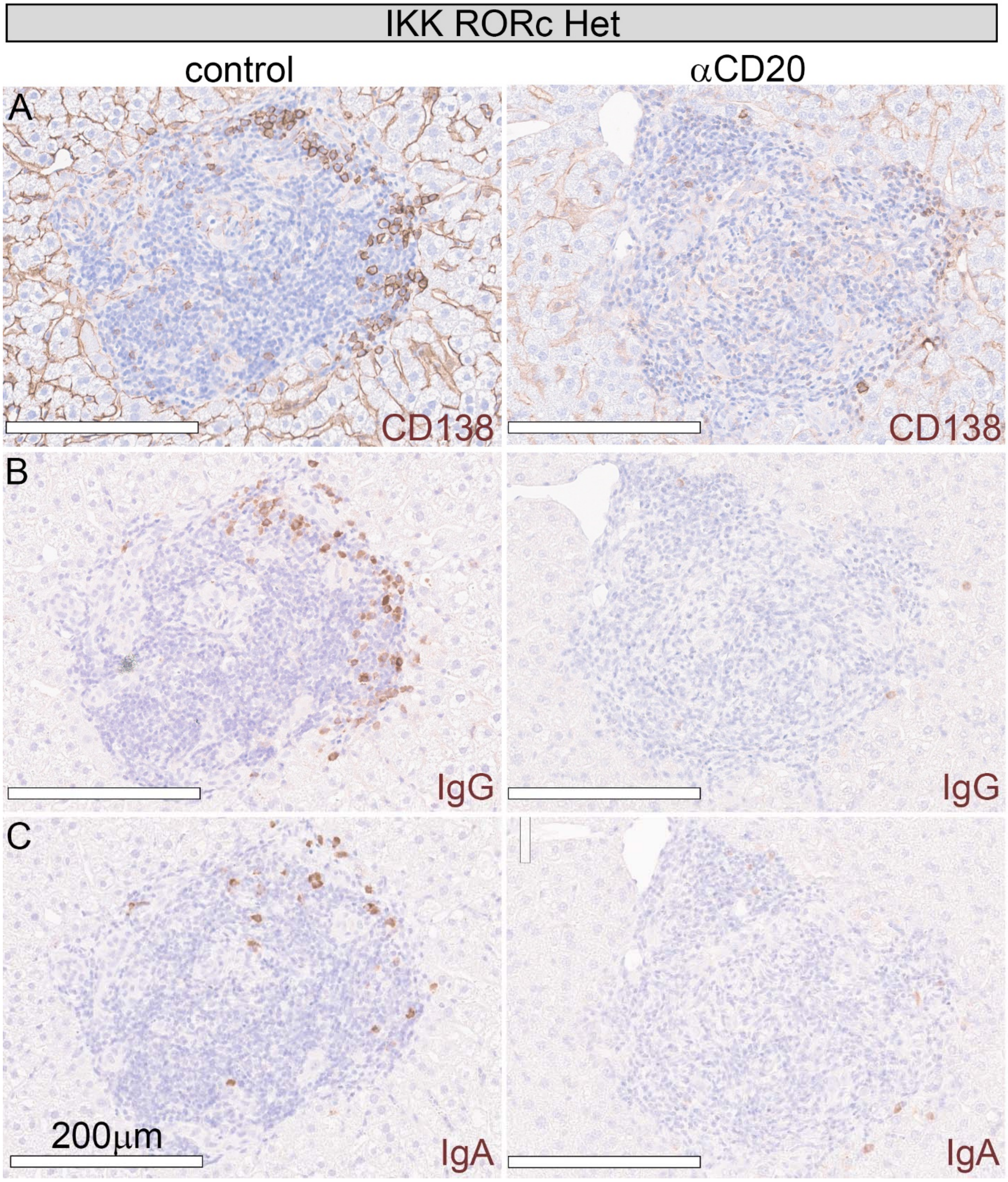

IKK RORc Het
control
αCD20
A
CD138
CD138
B
IgG
IgG
C
200μm
IgA
IgA

**Figure S20.**

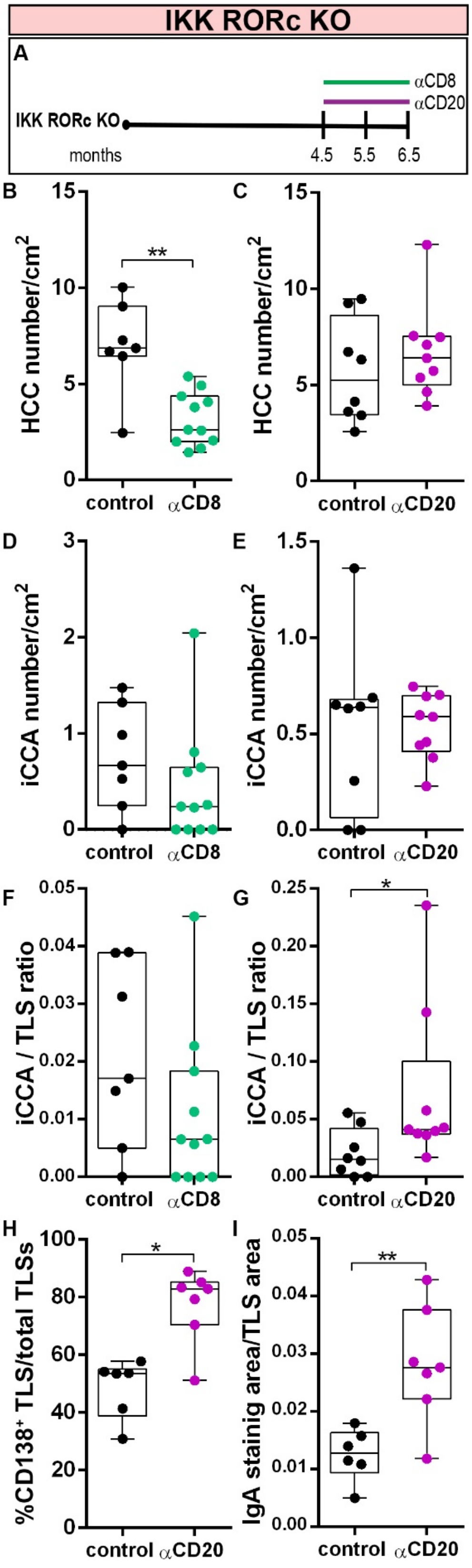
IKK RORc KO
A
αCD8
αCD20
IKK RORc KO
months
4.5
5.5
6.5
B
HCC number/cm2
**
control
αCD8
C
HCC number/cm2
control
αCD20
D
iCCA number/cm2
control
αCD8
E
iCCA number/cm2
control
αCD20
F
iCCA / TLS ratio
control
αCD8
G
iCCA / TLS ratio
*
control
αCD20
H
%CD138+ TLS/total TLSs
*
control
αCD20
I
IgA stainig area/TLS area
**
control
αCD20

**Figure S21.**

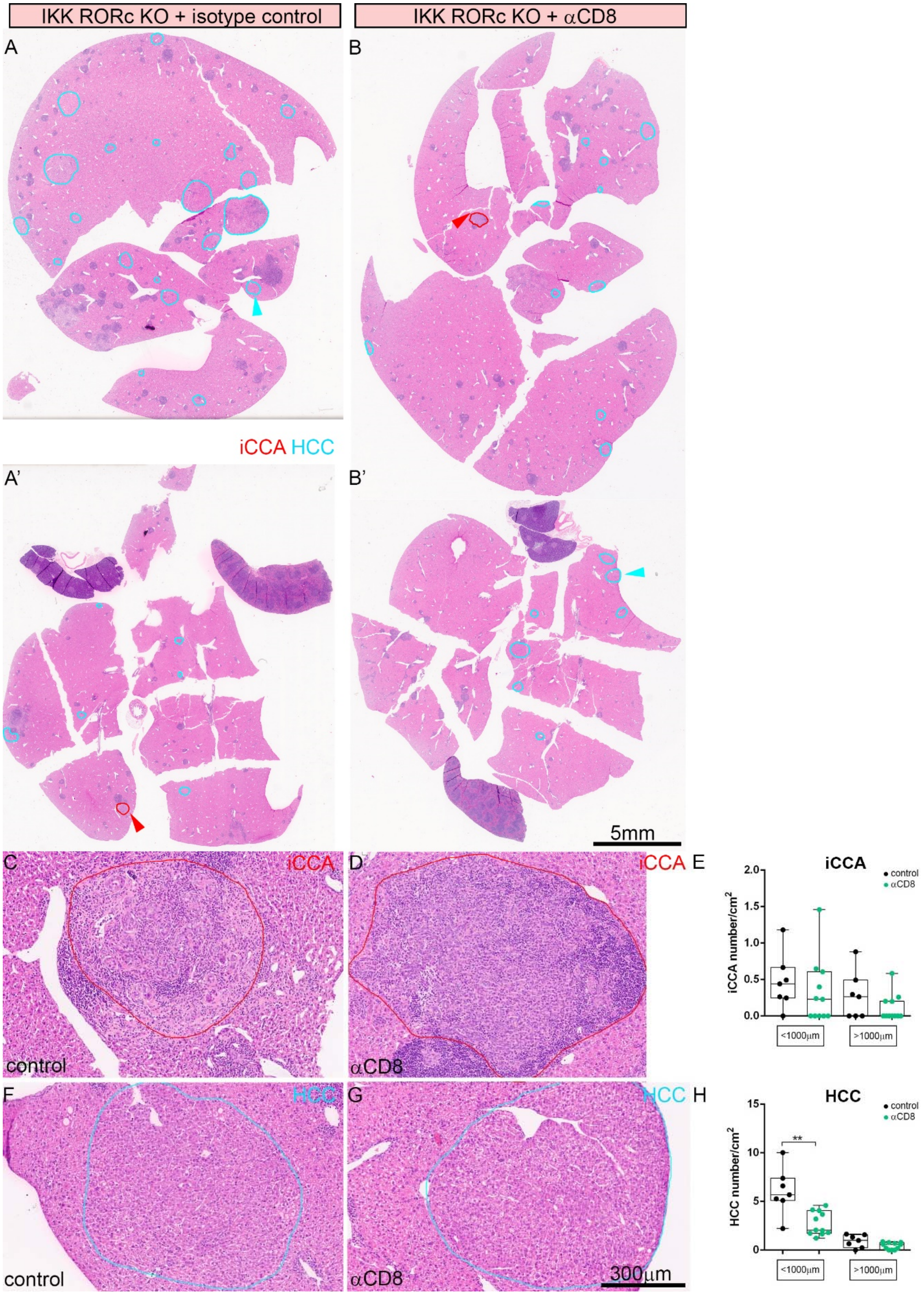
IKK RORc KO + isotype control
IKK RORc KO + αCD8
A
B
iCCA HCC
A'
B'
5mm
C
iCCA
D
iCCA
E
iCCA
control
αCD8
2.0
1.5
1.0
0.5
0.0
iCCA number/cm²
<1000μm
>1000μm
control
αCD8
F
HCC
G
HCC
H
HCC
control
αCD8
15
10
5
0
HCC number/cm²
**
<1000μm
>1000μm
300μm

**Figure S22.**

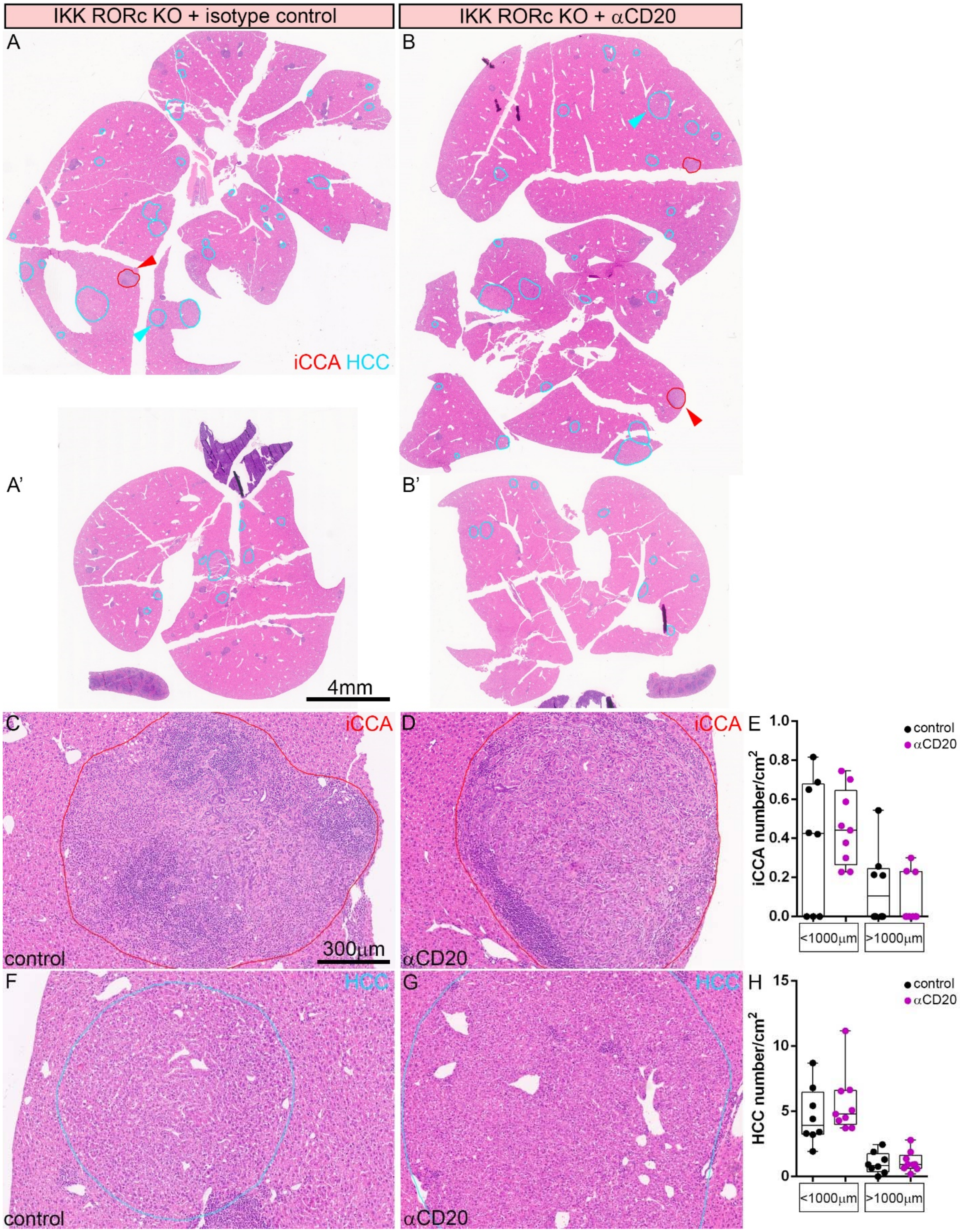

IKK RORc KO + isotype control
IKK RORc KO + αCD20
A
B
iCCA HCC
A'
B'
4mm
C
iCCA
D
iCCA
E
control
αCD20
iCCA number/cm²
1.0
0.8
0.6
0.4
0.2
0.0
<1000μm
>1000μm
control
300μm
αCD20
F
HCC
G
HCC
H
control
αCD20
HCC number/cm²
15
10
5
0
<1000μm
>1000μm
control
αCD20

**Figure S23.**

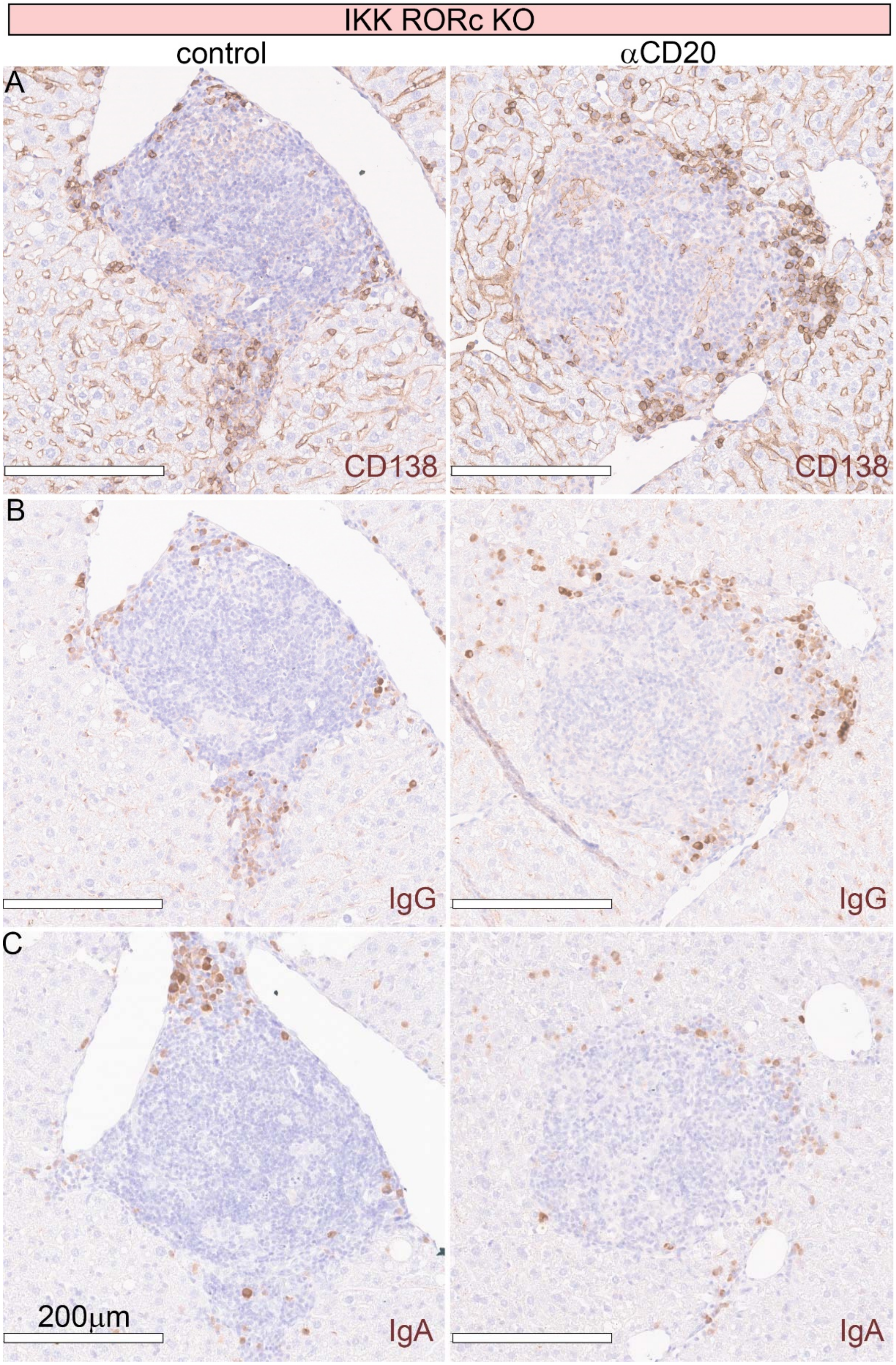

IKK RORc KO
control
αCD20
A
CD138
CD138
B
IgG
IgG
C
200μm
IgA
IgA
